# Expanded tRNA methyltransferase family member TRMT9B regulates synaptic growth and function

**DOI:** 10.1101/2022.12.30.522321

**Authors:** C.A. Hogan, S.J. Gratz, J.L. Dumouchel, A. Delgado, J.M. Lentini, K.R. Madhwani, R.S. Thakur, D. Fu, Kate M. O’Connor-Giles

## Abstract

Nervous system function relies on the formation and function of synaptic connections between neurons. Through a genetic screen in *Drosophila* for new conserved synaptic genes, we identified CG42261/Fid/ TRMT9B as a negative regulator of synaptogenesis. TRMT9B has been studied for its role as a tumor suppressor in multiple carcinomas and is one of two metazoan homologs of yeast tRNA methyltransferase 9 (Trm9), which methylates tRNA wobble uridines. Members of the expanded family of tRNA methyltransferases are increasingly being associated with neurological disorders and new biochemical functions. Interestingly, whereas Trm9 homolog ALKBH8/CG17807 is ubiquitously expressed, we find that TRMT9B is enriched in the nervous system, including at synapses. However, in the absence of animal models the role of TRMT9B in the nervous system has remained unknown. Here, we generated null alleles of *TRMT9B* and *ALKBH8*, and through liquid chromatography-mass spectrometry find that ALKBH8 is responsible for canonical tRNA wobble uridine methylation under basal conditions. In the nervous system, we find that TRMT9B negatively regulates synaptogenesis through a methyltransferase-dependent mechanism in agreement with our modeling studies. Finally, we find that neurotransmitter release is impaired in *TRMT9B* mutants. Our findings reveal a role for TRMT9B in regulating synapse formation and function, and highlight the importance of the expanded family of tRNA methyltransferases in the nervous system.

## Introduction

Nervous system function rests on communication between neurons at chemical synapses. Complex nervous systems are composed of thousands to billions of neurons connected through synapses in neural circuits that control behavior and thought. To achieve this functional organization, neurons must form synaptic connections in appropriate numbers and strength during development. Synapse number and strength are also substrates for the neuronal plasticity that underlies experience-dependent changes to the brain function (Citri and Malenka, 2008). Dysregulation of synapse formation and plasticity have been broadly linked to neurodevelopmental disorders, including autism spectrum disorder and other forms of intellectual disability (Guang et al., 2018; Zoghbi and Bear, 2012). While we have learned a great deal about a small subset of genes underlying neurodevelopmental disorders, a recent study found that the majority of genes in the human genome are understudied – including those associated with human disorders (Stoeger et al., 2018). With the increasingly broad application of clinical sequencing in individuals with neurodevelopmental delay, new genes whose function in the nervous system remains unknown are being uncovered. A lack of foundational understanding of synaptic gene function is a significant obstacle for translating clinical identification of causative variants into effective treatments for neurodevelopmental disorders.

To uncover conserved, uncharacterized genes with roles in the formation of functional synapses, we conducted a genetic screen in *Drosophila* for regulators of synaptic growth at the well-characterized glutamatergic neuromuscular junction (NMJ). Computed Gene 42261 (CG42261) emerged from our screen as a negative regulator of synaptic growth. CG42261, which was also identified in an RNAi screen for regulators of thermal nociception and named *fire dancer* (Honjo et al., 2016), encodes the *Drosophila* homolog of human TRMT9B. TRMT9B has been studied for its role as a tumor suppressor downregulated in colorectal, breast, bladder, cervical, ovarian and testicular carcinomas (Begley et al., 2013; Chen et al., 2017; Flanagan et al., 2004; Wang et al., 2018). TRMT9B is one of two metazoan homologs of yeast tRNA methyltransferase 9 (Trm9), which methylates tRNA wobble uridines (Begley et al., 2007; Begley et al., 2013; Fu et al., 2010; Kalhor and Clarke, 2003; Songe-Moller et al., 2010). tRNAs bridge mRNA decoding and protein synthesis, and are among the most heavily post-transcriptionally modified RNAs (Boccaletto et al., 2022). These modifications affect the secondary structure and stability of tRNAs as well as the efficiency and fidelity of codon interactions (Agris et al., 2017). tRNA wobble uridines are unique in that they are nearly always modified across organisms through complex enzymatic pathways that remain incompletely understood (Chen et al., 2011; Schaffrath and Leidel, 2017). Trm9 homolog ALKBH8 has a demonstrated role in catalyzing tRNA wobble uridine methylation in mammals (Fu et al., 2010a; Songe-Moller et al., 2010), whereas TRMT9B’s role has remained unknown despite years of study. Intriguingly, an emerging body of work demonstrates that enzymes in the tRNA methytransferase family have evolved to methylate diverse substrates, including rRNAs, mRNAs and proteins (Abbasi-Moheb et al., 2012; Chellamuthu and Gray, 2020; Chen et al., 2011; Xu et al., 2017). Methylation of each of these substrates provides a reversible level of regulation that modulates biological processes ranging from transcription to splicing to translation to protein interactions (Greenberg and Bourc’his, 2019; Murn and Shi, 2017; Zhou et al., 2020).

Here, we detail the first animal model of *TRMT9B* to investigate its biological role in the nervous system. Using endogenously tagged lines, we find that TRMT9B is expressed in neuronal, glial and muscle cell bodies, and at synapses. At the NMJ, TRMT9B functions post-synaptically to attenuate synaptic growth and promote neurotransmitter release. Through quantitative RNA mass spectrometry, we determined that TRMT9B is dispensable for baseline tRNA wobble uridine methylation. Our modeling studies support TRMT9B’s role as a Class I SAM-dependent methyltransferase, and we demonstrate that TRMT9B regulates synapse development through a methyltransferase-dependent mechanism. Together our findings highlight the expanding roles of the tRNA methyltransferase family in animals and reveal a critical requirement for TRMT9B-dependent methylation in nervous system development.

## Results

### TRMT9B negatively regulates synaptic growth

As part of an ongoing effort to identify and characterize new regulators of synapse formation and function, we used ModENCODE developmental RNA-Seq data to identify uncharacterized, conserved neuronal genes whose expression in *Drosophila* corresponds to periods of synaptogenesis (Graveley et al., 2011). We then knocked down top candidate genes using existing RNAi lines (Dietzl et al., 2007; Hu et al., 2017; Ni et al., 2011) and screened for altered synapse number at glutamatergic motor synapses. Knockdown of CG42261, which encodes one of two *Drosophila* homologs of yeast wobble uridine methyltransferase Trm9, resulted in significant ectopic synapse formation. CG42261 was also identified in an RNAi screen for genes required for thermal nociception and named *fire dancer (fid)* based on observed deficits (Honjo et al., 2016). Here, we follow recently established nomenclature for members of the tRNA methyltransferase family and refer to CG42261/Fid as TRMT9B (Tweedie et al., 2021).

To confirm our findings, we used CRISPR-based gene editing to generate two independent null alleles: (1) *TRMT9B*^*KO*^, in which all coding exons are removed and replaced with a visible marker and attP landing site and (2) *TRMT9B*^*HA+IC*^, in which an HA peptide tag and a cassette encoding a visible marker flanked by piggyBac inverted terminal repeat sequences is inserted immediately following the sole *TRMT9B* translational start site (see Materials and methods). The visible marker cassette disrupts the open reading frame to generate a null allele. Its subsequent removal by piggyBac transposase yields an in-frame epitope-tagged allele (Bruckner et al., 2017).

At well-characterized *Drosophila* larval NMJs, individual motorneurons form tens to hundreds of glutamatergic synapses with postsynaptic muscle targets in a stereotyped manner. We quantified synaptic growth by counting synaptic boutons at the highly stereotyped NMJ 4 formed between motorneuron 4-1b and muscle 4. Consistent with our screening results, we observed a significant increase in bouton number in homozygous *TRMT9B*^*KO*^, homozygous *TRMT9B*^*HA+IC*^, and heteroallelic *TRMT9B*^*KO/HA+IC*^ mutants (Fig. 1A-D,F). All three TRMT9B allelic combinations exhibited a 30-35% increase in bouton number relative to control. We observe similar synaptic overgrowth in *TRMT9B*^*KO*^ over a deficiency, and in a line with an artificial stop codon (*TRMT9B*^*MiMIC-STOP*^) that is partially restored to wild type when one copy is replaced with a GFP codon (*TRMT9B*^*MiMIC-GFP*^, Fig. S1; Nagarkar-Jaiswal et al., 2015; Venken et al., 2011).

**Figure 1.**
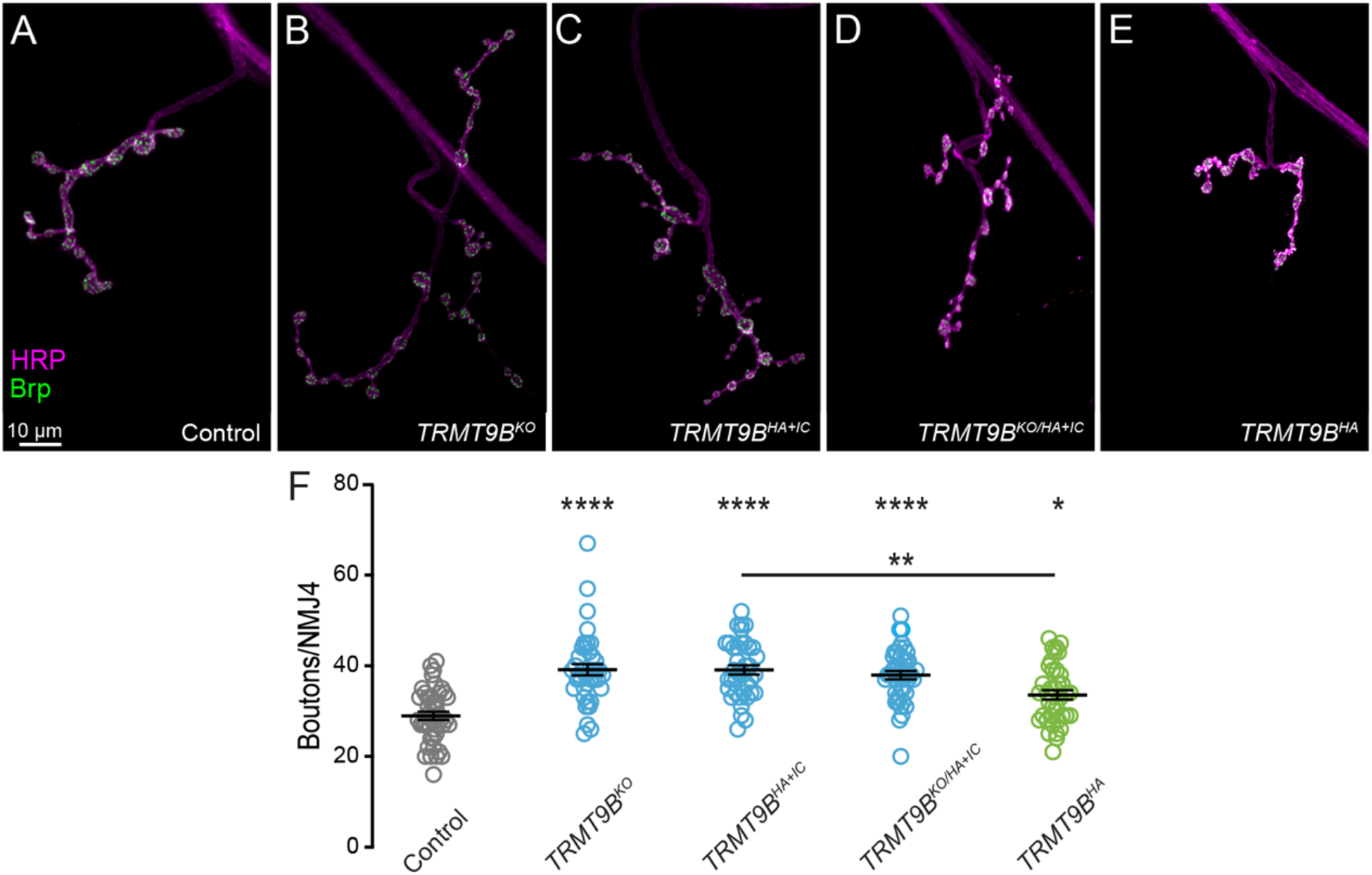
TRMT9B attenuates synaptic growth. (**A–F**) Representative confocal images (A-E) and bouton number per NMJ 4 (**F**) in the indicated genotypes. Three combinations of *TRMT9B* null alleles exhibit synaptic overgrown relative to control, and restoration of *TRTM9B* under endogenous control significantly rescues synaptic overgrowth. Kruskal-Wallis test followed by Dunn’s multiple comparisons test. Error bars represent s.e.m.

To further confirm loss of *TRMT9B* as the cause of synaptic overgrowth, we restored gene function under endogenous control by removing the interfering cassette in *TRMT9B*^*HA+IC*^ to generate N-terminally tagged *TRMT9B*^*HA*^. Synapse number at *TRMT9B*^*HA*^ NMJs was restored to near control levels (Fig. 1E,F).

### TRMT9B functions postsynaptically to attenuate synaptic growth

Synapse formation involves a complex interplay between pre- and postsynaptic cells. Our endogenous rescue system offers the advantage of driving expression at biological levels, but does not allow spatial control of gene expression. To determine the site of TRMT9B function in regulating motor synapse formation, we conducted cell-specific rescue using *C155-Gal4* and *24B-Gal4* to drive expression of a full-length *TRMT9B* transgene in presynaptic neurons and postsynaptic muscle, respectively. We found that muscle, but not neuronal, expression of *TRMT9B* fully rescues synaptic growth at the NMJ (Fig. 2). Thus, TRMT9B acts post-synaptically to attenuate synaptic growth.

**Figure 2.**
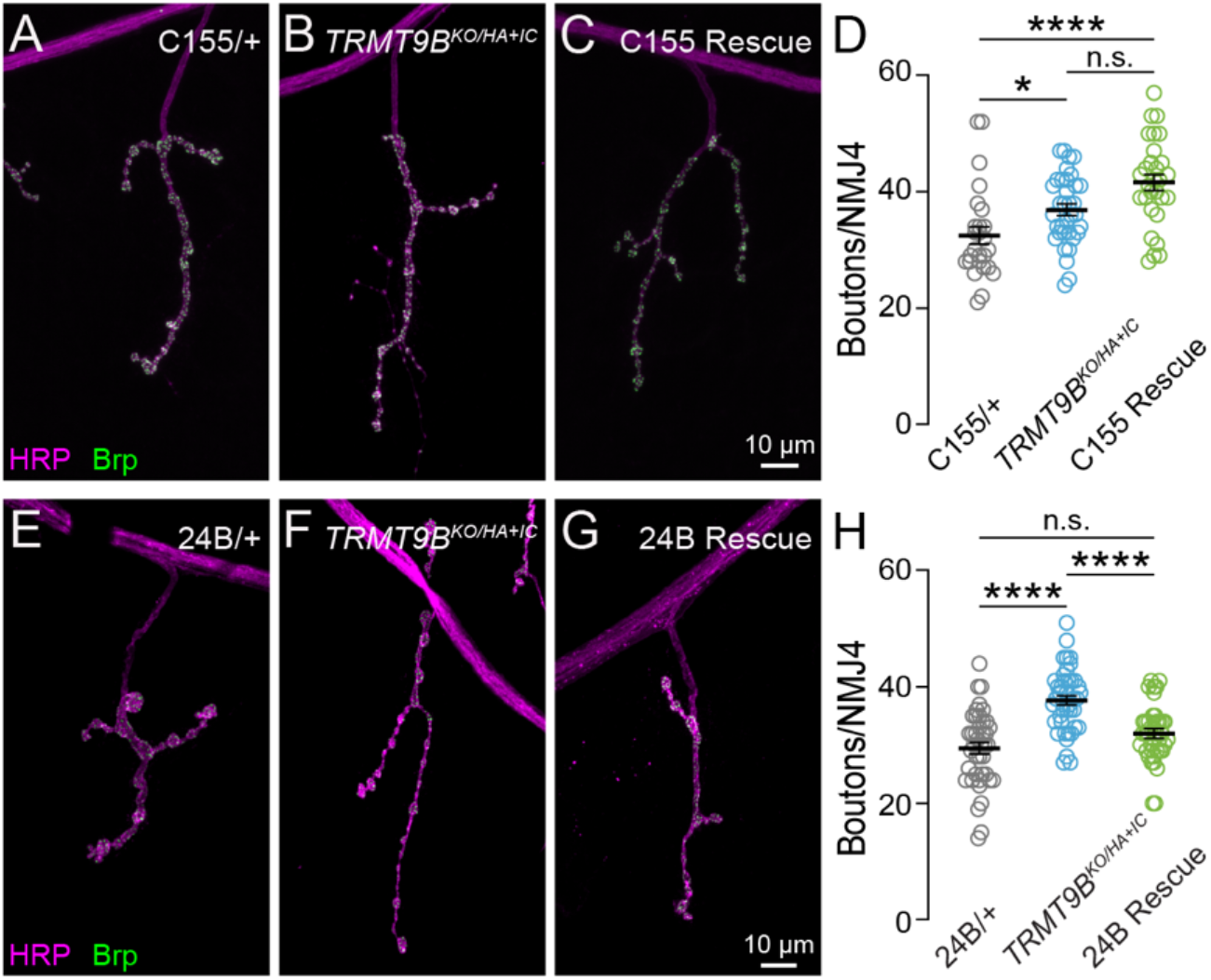
TRMT9B functions postsynaptically to attenuate synaptic growth. (**A–H**) Representative confocal images of NMJ 4 in indicated genotypes with corresponding bouton number per NMJ. **(A-D)** Presynaptic restoration of TRMT9B does not rescue synaptic overgrowth **(E-H)** In contrast, postsynaptic restoration of *TRMT9B* fully rescues synaptic overgrowth. Kruskal-Wallis test followed by Dunn’s multiple comparisons test. Error bars represent s.e.m.

### Wobble uridine methylation in the absence of TRMT9B

TRMT9B and its paralog ALKBH8 are homologous to yeast Trm9, yet it has remained unknown if both paralogs carry out the canonical role of methylating wobble uridines. Of the two metazoan homologs, TRMT9B has a domain structure more similar to Trm9 with a well-conserved Class I SAM-dependent methyltransferase domain (Fig. 3A). In contrast, ALKBH8, in addition to a conserved methyltransferase domain, contains an RNA-binding motif and a 2OG-Fe (II) oxygenase domain that catalyzes an animal-specific tRNA modification (Fig. 3A; (Fu et al., 2010b; van den Born et al., 2011). Mouse and human ALKBH8 have been shown to methylate tRNA wobble uridines (Fu et al., 2010a; Songe-Moller et al., 2010), whereas TRMT9B’s biochemical function has remained unknown in the absence of animal models. To investigate the role of the two *Drosophila* Trm9 paralogs in tRNA methylation, we generated a null allele of *Drosophila ALKBH*8, currently identified as CG17807. To our knowledge, this is the first animal model in which both paralogs have been knocked out, allowing us to discern the independent molecular function of each protein.

**Figure 3.**
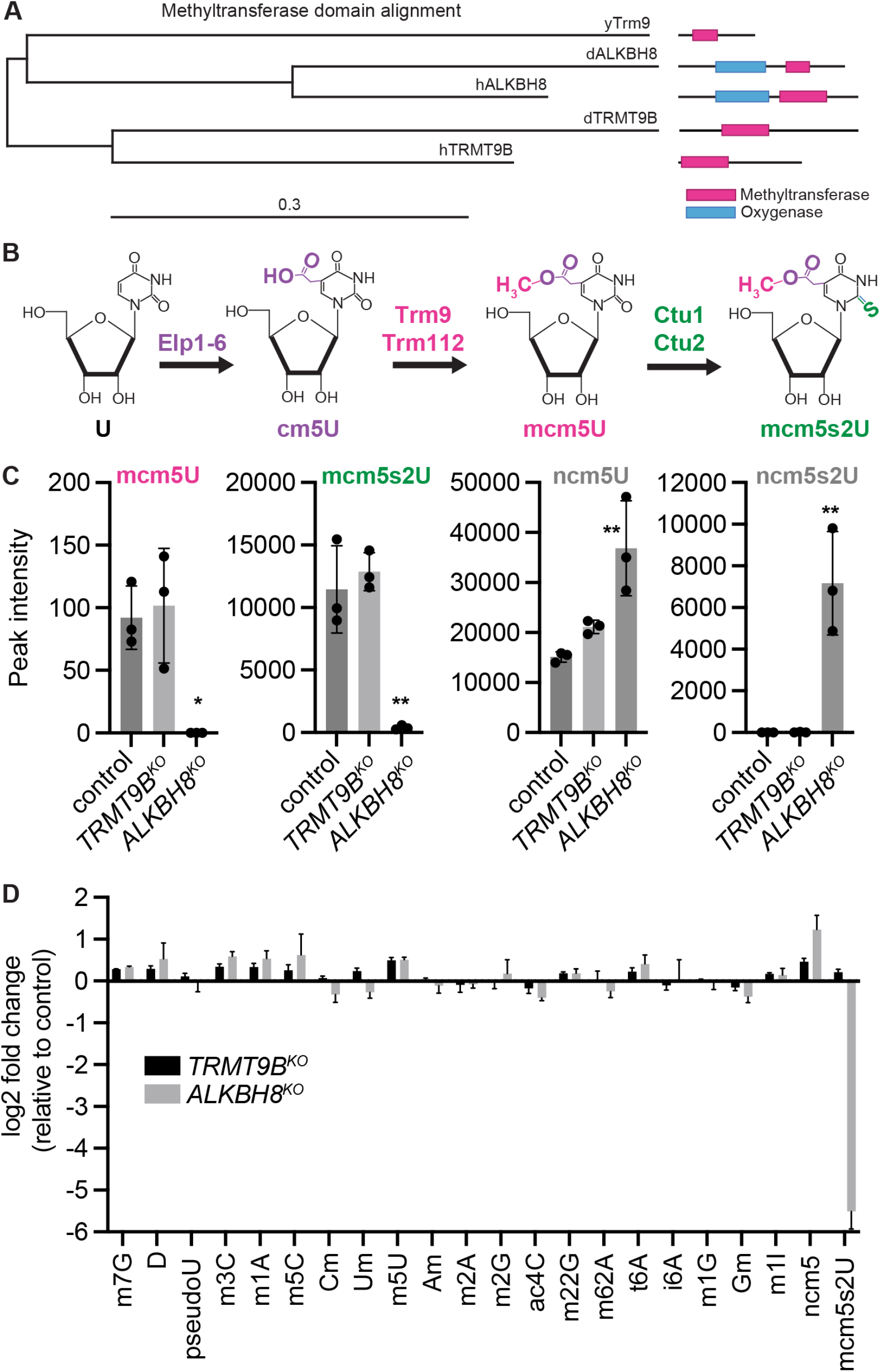
TRMT9B is not required for tRNA wobble uridine methylation. (**A**) Phylogenetic tree of yeast Trm9, fly and human TRMT9B, and fly and human ALKBH8 determined by maximum likelihood from alignments or their methyltransferase domains. (**B**) Yeast wobble uridine methylation pathway. (**C**) Peak intensity areas of modified uridines normalized to the canonical nucleosides A, U, G, and C in total RNA isolated from control, *TRMT9B*^*KO*^ and *ALKBH8*^*KO*^ heads. (**D**) Log2 fold change in the levels of the indicated tRNA modification between *TRMT9B*^*KO*^ and *ALKBH8*^*KO*^ heads relative to control strains. Samples were measured in triplicate. Error bars represent standard deviation.

Most of our understanding of tRNA wobble uridine modification comes from studies in yeast, where posttranscriptional modification modulates tRNA interactions with cognate and wobble codons (Schaffrath and Leidel, 2017). In one branch of the pathway, the Elongator complex is thought to act first to generate 5-carbonylmethyluridine (cm5U, Fig. 3B, purple).Trm9 then acts in concert with obligate cofactor Trm112 to add a second methyl group and generate 5-methoxycarbonylmethyluridine (mcm5U, Fig. 3B, pink). Further modification by the Ncs2-Ncs6 thiolase complex generates a terminal mcm5s2U modification (Fig. 3B, green). In a second branch, wobble uridines in certain tRNAs are converted to the amide modification ncm5U by an unknown enzyme (Johansson et al., 2008).

To analyze tRNA wobble uridine modification in our null alleles, we conducted quantitative liquid chromatography-mass spectrometry (LC-MS). Briefly, total RNA isolated from heads of mated male and female control, *TRMT9B*^*KO*^ and *ALKBH8*^*KO*^ flies was nuclease digested and dephosphorylated to generate individual ribonucleosides followed by LC-MS analysis of nucleoside modifications (Cai et al., 2015; Dewe et al., 2017; Fu et al., 2010a). We found that the mcm5U and mcm5s2U modifications, which are tRNA specific and in yeast depend on Trm9, are present in nucleosides isolated from control and *TRMT9B*^*KO*^ flies, but reduced to near background levels in *ALKBH8*^*KO*^ flies (Fig. 3C).

Notably, we detected an increase in the accumulation of ncm5U and ncm5S2U in *ALKBH8*^*KO*^ flies, but not control or *TRMT9B* flies (Fig. 3C, D). These findings are consistent with studies in yeast showing that loss of Trm9 leads to loss of mcm5U and mcm5s2U concomitant with the accumulation of amide intermediates (Chen et al., 2011). Based on these findings, we conclude that ALKBH8 methylates tRNA wobble uridines independently of TRMT9B under basal conditions.

We next explored the possibility that TRMT9B catalyzes a different tRNA methylation – a possibility hinted at in previous overexpression studies (Begley et al., 2013). Specifically, we investigated a panel of well-characterized tRNA modifications at multiple tRNA sites, the majority involving methylation.Further confirming our results above, we found that mcm5s2U was decreased to near background levels in *ALKBH8*^*KO*^ flies while no major change in mcm5s2U was detected in *TRMT9B*^*KO*^ flies (Fig. 3D). For the remaining modifications with levels within a quantifiable range, we observed a slight variation in levels for a subset of nucleosides isolated from the *ALKBH8*^*KO*^ flies when compared to control flies, suggesting that loss of ALKBH8-catalyzed wobble uridine modifications could impact other modifications in tRNA. In contrast, no modification exhibited more than a 2-fold change in the *TRMT9B*^*KO*^ compared to control. While our findings do not rule out the possibility that TRMT9B plays a conditional role in tRNA wobble uridine methylation or methylates a small subset of tRNAs not detectable by bulk analysis, these observations indicate that ALKBH8 is the primary paralog carrying out the canonical Trm9 tRNA wobble uridine methyltransferase role and suggest a new role for TRMT9B.

### TRMT9B has a structurally conserved methyltransferase domain

This finding raises the question of whether TRMT9B has maintained function as a methyltransferase of evolved a non-enzymatic role. To investigate TRMT9B’s role as a methyltransferase, we first aligned the predicted methyltransferase domains of yeast Trm9 with *Drosophila* and human TRMT9B (Fig. 4A). Although methyltransferases are not generally well conserved at the level of primary sequence (Martin and McMillan, 2002), we observe conservation throughout the methyltransferase domain, including residues known to be important for methyltransferase function. The defining sequence features of Class I SAM-dependent methyltransferase are (1) an acidic residue at the end of the first beta strand and (2) a GXGXGX SAM-binding motif – both of which are critical for SAM binding and conserved in yeast Trm9 and fly and human TRMT9B (Fig. 4A, blue boxed residues; Kozbial and Mushegian, 2005; Martin and McMillan, 2002). Additional acidic residues in beta strands 2, 3 and 4 and glycines in beta strand 5 that are highly conserved across Class I SAM-dependent methyltransferases are also present in fly and human TRMT9B (Fig. 4A, blue residues; Kozbial and Mushegian, 2005). We next assessed residues found to be critical for SAM binding and enzymatic activity in a comprehensive mutational analysis of yeast Trm9 (Létoquart et al., 2015), and found that these residues are also highly conserved in *Drosophila* and human TRMT9B and ALKBH8 (Fig. 4A, black and red boxed residues). Together, these observations are consistent with the model that both paralogs retain methyltransferase function.

**Figure 4.**
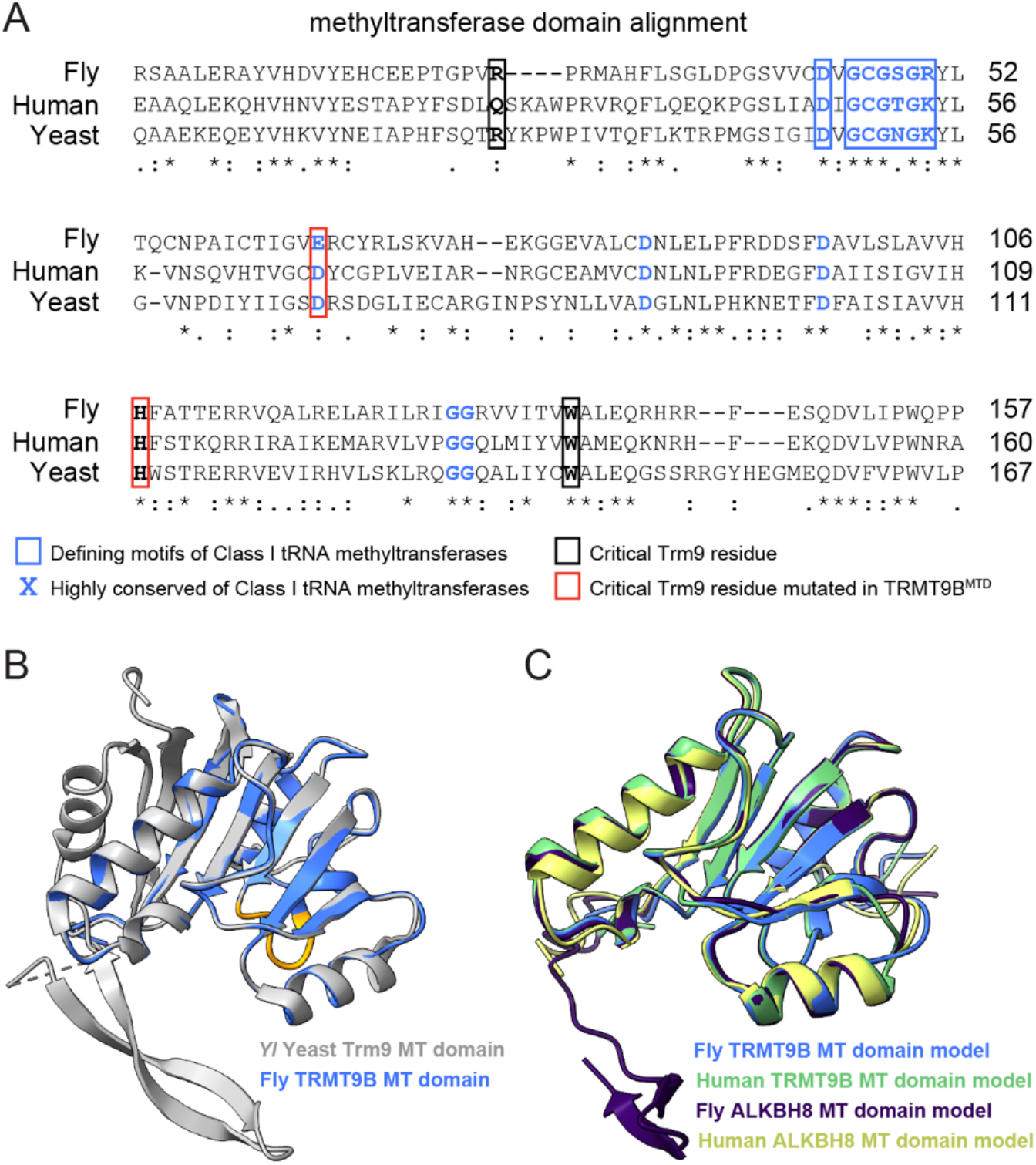
TRMT9B encodes a conserved methyltransferase. (**A**) Sequence alignment of Yeast Trm9, *Drosophila* TRMT9B and human TRMT9B methyltransferase domains. Class I methyltransferase defining sequences are blue and boxed. Additional residues highly conserved across class I methyltransferases are indicated in blue. Critical residues identified in yeast by Letoquart *et*.*al*. 2015 are boxed in black or red, with red indicating the amino acids mutated in our methyltransferase-dead transgene. Identical amino acids are marked with asterisks, conserved substitutions are marked with two dots, and semi-conserved substitutions with single dots. (**B**) Structural homology model of the *Drosophila* TRMT9B methyltransferase domain (blue) with yeast Trm9 structure (gray, PBD ID: 5CM2 chain Z). The class I methyltransferase GxGxGx motif (orange) and acidic SAM binding residue (D72 in yeast and E189 in fly) are in conserved positions across species. (**C**) Extension of the homology model shows conserved secondary structure between *Drosophila* (blue) and human (green) TRMT9B and *Drosophila* (purple) and human ALKBH8 (yellow) methyltransferase domains.

To investigate secondary structure, we generated structural homology models of the methyltransferase domains of *Drosophila* and human TRMT9B and ALKBH8. For an unbiased approach, we used Modeller to model the methyltransferase domains against known crystal structures reported in the RCSB Protein Data Bank (Berman et al., 2000; Pieper et al., 2014) and identified Trm9 as a match for each enzyme, lending further support for the hypothesis that both TRMT9B and ALKBH8 maintain methyltransferase function. To evaluate structural conservation, we used Chimera to compare our *Drosophila* TRMT9B methyltransferase domain model to the yeast Trm9 structure determined by X-ray diffraction of *Yarrowia lipolytica* Trm9 and its obligate co-factor Trm112 (Fig. 4B; Létoquart et al., 2015). Class I methyltransferases form a seven-stranded beta sheet flanked by alpha helices. In our model, this folded structure is maintained and key residues are superimposed (Fig. 4B, orange ribbon). We observed similar structural conservation between yeast Trm9 and our model of human TRMT9B (Fig. S2). We next overlaid our models of TRMT9B with *Drosophila* and human ALKBH8, and observed significant structural conservation between the four methyltransferase domains (Fig. 4C). Finally, we assessed the predicted structure of the *Drosophila* TRMT9B methyltransferase domain using AlphaFold (Jumper et al., 2021; Varadi et al., 2022). Strong alignment was observed between the AlphaFold predicted model, the *Drosophila* TRMT9B model predicted using Modeller, and the yeast TRM9 crystal structure (Fig. S3). Thus, multiple independent *in silico* modeling approaches support structural conservation of TRMT9B’s methyltransferase domain across species, suggesting TRMT9B functions as a methyltransferase in flies and humans.

### TRMT9B regulation of synaptic growth requires methyltransferase function

To experimentally assess whether TRMT9B functions as a methyltransferase in the nervous system, we generated a methyltransferase-dead rescue construct for expression under GAL4 control and compared its ability to rescue synaptic growth deficits in *TRMT9B* mutants to wild-type *TRMT9B*. To disrupt methyltransferase function, we replaced two critical residues identified by Letoquart and colleagues with alanines (See Fig. 3A, red boxed residues; Létoquart et al., 2015). We modeled the effect of these mutations on protein structure and found that secondary protein structure remains intact, consistent with prior studies (Fig. S4; Létoquart et al., 2015). We integrated wild-type (*UAS-TRMT9B*) and methyltransferase-dead (*UAS-TRMT9B*^*MTD*^) transgenes at the same genomic site and expressed each transgene under the control of *24B-GAL4*. While expression of wild-type *TRMT9B* fully rescued synaptic overgrowth as observed above, expression of *TRMT9B*^*MTD*^ failed to rescue (Fig. 5), consistent with the conclusion that methyltransferase activity is required for TRMT9B’s role in regulating synaptic growth. To confirm proper localization of methyltransferase-dead TRMT9B, we generated V5-tagged versions of the wild-type and methyltransferase-dead transgenes and confirmed similar localization patterns when expressed in muscle (Fig. S5). Consistent with our sequence and structural analyses, these data indicate that TRMT9B regulates synaptic growth through a methyltransferase-dependent mechanism.

**Figure 5.**
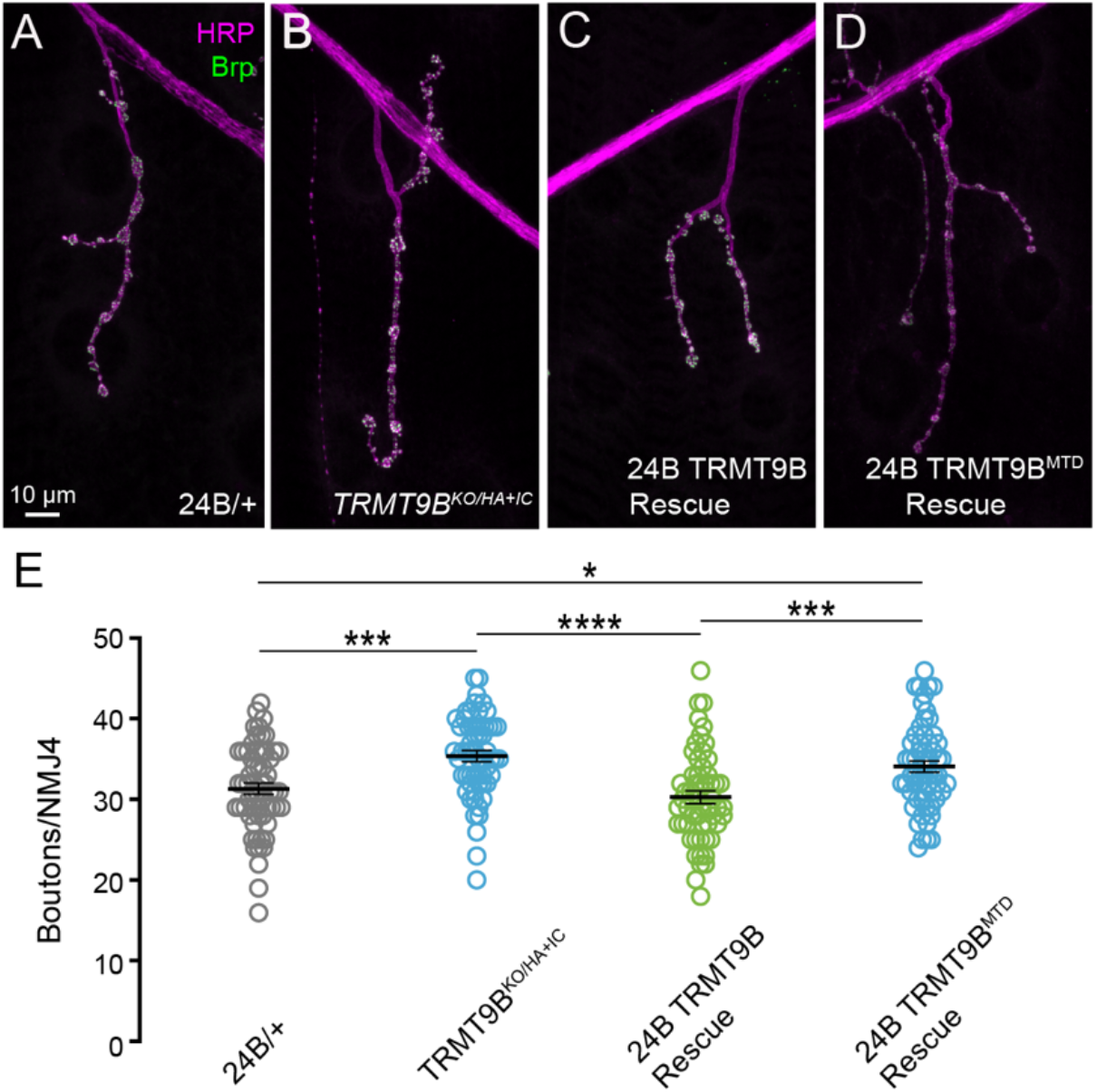
TRMT9B regulation of synaptic growth requires methyltransferase function. (**A– E**) Representative confocal images (**A-D**) and bouton number per NMJ 4 (**E**) in the indicated genotypes. *24B*-Gal4-driven expression of wild-type *TRTM9B* significantly rescues synaptic overgrowth. In contrast, 24B-Gal4-driven expression of methyltransferase-dead *TRTM9B* fails to rescue synaptic overgrowth. ANOVA followed by Tukey’s multiple comparisons test. Error bars represent s.e.m.

### TRMT9B is expressed in neurons, glia and muscle

In contrast with ALKBH8, which is ubiquitously expressed across species, *TRMT9B* is enriched in *Drosophila* and mammalian nervous systems (Brown et al., 2014; GTEx Consortium, 2015). To investigate TRMT9B protein expression in the nervous system and, specifically, at the NMJ, we turned to our endogenously tagged alleles. In N-terminally tagged *TRMT9B*^*sfGFP*^, we observe expression in neuronal cell bodies (Fig. 6A) and the synaptic neuropil of the larval ventral ganglion as indicated by colocalization with the active zone protein Brp (Fig. 6B). The synaptic localization is consistent with previous observations from a large-scale MiMIC protein trap collection in which TRMT9B is internally tagged prior to the methyltransferase domain (Fig. S6A; Nagarkar-Jaiswal et al., 2015). We also observe a similar expression pattern, albeit at lower levels, in *TRMT9B*^*HA*^, which rescues synaptic growth and function (Fig. S6B; See Figs 1 and 6), indicating that tagged alleles maintain function and localize similarly in the nervous system. To determine which cell types express TRMT9B in the nervous system, we conducted colocalization studies with the neuronal marker Elav and glial marker Repo. We found that TRMT9B is strongly expressed in neuronal as well as glia cell bodies (Fig. 6C). In muscle where TRMT9B attenuates synaptic growth, TRMT9B is broadly cytoplasmic with peri-nuclear accumulation as observed with our V5-tagged transgene (Fig. 6D, See Fig. S5). Finally, we observe TRMT9B at the NMJ where it primarily colocalizes presynaptically with HRP, although we cannot rule out expression at the postsynaptic density (Fig. 6D). The subcellular and synaptic localization patterns of TRMT9B in neurons, glia and muscle raise the intriguing possibility of a role for TRMT9B in methylating multiple substrates at distinct subcellular locations, including at synapses.

**Figure 6.**
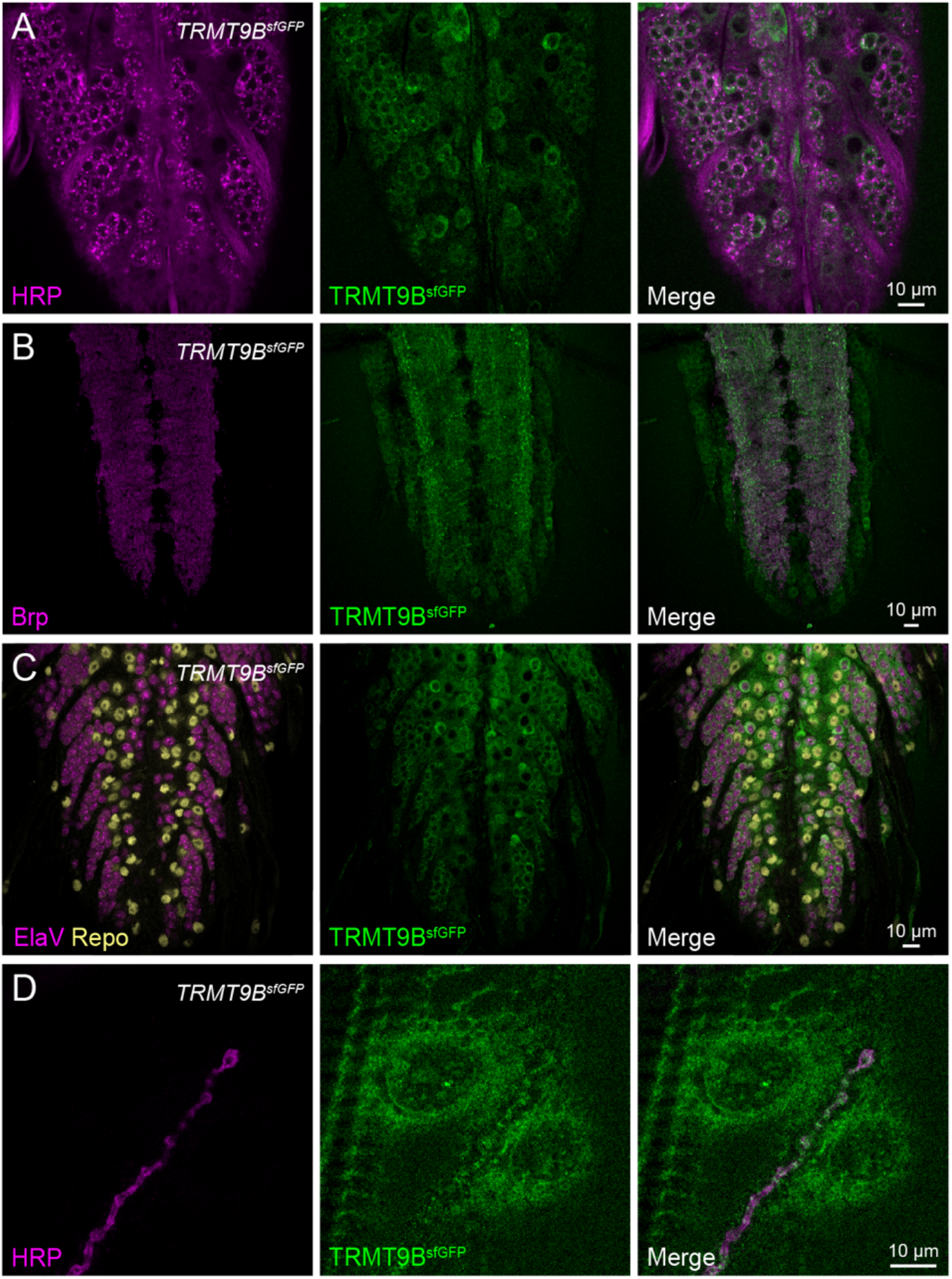
TRMT9B is expressed in neuronal, glial and muscle cell bodies and at synapses. (**A**) Confocal Z-projections of a *TRMT9B*^*sfGFP*^ larval ventral ganglion co-labeled with antibodies against GFP (green) and the neuronal membrane marker HRP (magenta) showing expression in cell bodies. (**B**) Confocal Z-projections of a *TRMT9B*^*sfGFP*^ larval ventral ganglion co-labeled with antibodies against GFP (green) and the synaptic marker Brp (magenta) showing expression in the synaptic neuropil. (**C**) Confocal Z-projections of *TRMT9B*^*sfGFP*^ larval ventral ganglia co-labeled with antibodies against GFP (green) and neuronal marker Elav (magenta) or glial marker Repo (yellow) indicates expression in neurons and glia. (**D**) Confocal Z-projections of *TRMT9B*^*sfGFP*^ NMJ/muscle co-labeled with antibodies against GFP (green) and neuronal membrane marker HRP (magenta) reveals expression in boutons and muscle with perinuclear accumulation.

### TRMT9B promotes neurotransmitter release

To investigate additional roles for TRMT9B in the nervous system, we assessed neurotransmission at *TRMT9B* null NMJs. To assess neurotransmission in our mutants, we measured spontaneous and evoked synaptic potentials through current clamp recordings at muscle 6, which is innervated by one type Ib and one type Is motorneuron. We observed significantly reduced excitatory junction potentials (EJPs) together with normal miniature EJP (mEJP) amplitude (quantal size) in *TRMT9B* null mutants (Fig. 7A-E). These findings indicate a 51% decrease in neurotransmitter release (quantal content; Fig. 7F). Restoration of endogenous *TRMT9B* function through removal of the *TRMT9B*^*HA+IC*^ interfering cassette fully rescued neurotransmitter release (Fig. 7A-F, *TRMT9B*^*HA*^), confirming that loss of *TRMT9B* underlies the deficit in synaptic function. To assess where TRMT9B function is required for neurotransmission, we conducted cell-specific rescue using *C155-Gal4* and *24B-Gal4* to drive expression of full-length TRMT9B pre- and postsynaptically, respectively. As with synaptic growth, we found that postsynaptic, but not presynaptic, expression of TRMT9B fully rescues neurotransmitter release (Fig. S7).

**Figure 7.**
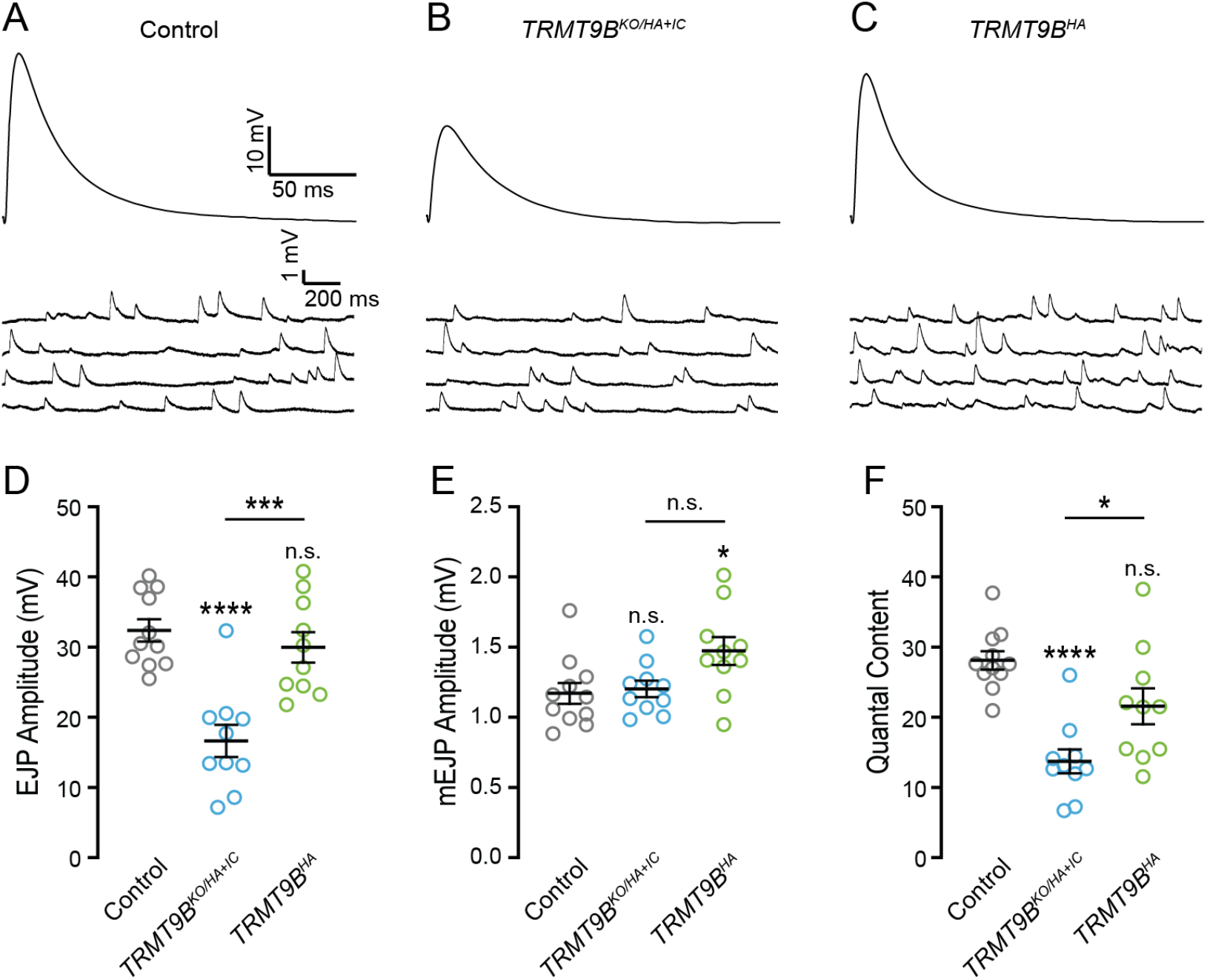
TRMT9B promotes neurotransmitter release. (**A–C**) Representative traces of EJPs and mEJPs in indicated genotypes. Stimulus artifacts have been removed from representative EJPs for clarity. (**D-F**) Average mEJP amplitude (**D**), EJP amplitude (**E**) and quantal content (**F**) for the indicated genotypes. mEJP amplitude is slightly elevated in *TRMT9B*^*HA*^ relative to control. EJP amplitude and quantal content are significantly diminished in *TRMT9B*^*KO/HA+IC*^ and rescued by restoration of *TRMT9B* expression. ANOVA followed by Tukey’s test. Error bars represent s.e.m.

Given that synaptic bouton number is increased in *TRMT9B* mutants (See Fig. 1), we reasoned that decreased neurotransmitter release could be due to the formation of fewer synapses per bouton or diminished function at individual synapses. To distinguish these possibilities, we quantified individual synapses in control and *TRMT9B* NMJs with the presynaptic marker Brp. We found that the number of synapses per bouton is decreased by 15% at *TRMT9B*^*KO*^ NMJs (9.43 ± 0.41 vs. 7.97 ± 0.33 synapses per bouton, p=0.013, Student’s t test). Combined with the 35% increase in bouton number, this yields a >10% increase in total synapse number at the NMJ, indicating that decreased synapse number does not underlie the deficit in neurotransmitter release and suggesting instead that individual *TRMT9B* synapses are functionally impaired. Thus, TRMT9B regulates both synapse number and the formation of functional synapses – defining novel roles in the nervous system for this member of the expanded family of tRNA methyltransferases.

## Discussion

Through a genetic screen to identify conserved regulators of synaptogenesis, we discovered a role for expanded tRNA methyltransferase family member TRMT9B in nervous system development. While TRMT9B has previously been associated with a number of non-neural carcinomas, its role in the nervous system where it is enriched has not previously been studied. Through the generation of the first animal model, we have found that TRMT9B acts postsynaptically to regulate synapse formation and function. Our biochemical, modeling, and *in vivo* studies indicate that TRMT9B functions through a methyltransferase-dependent mechanism and may have evolved in animals to methylate a novel substrate – highlighting both the expanding functions of tRNA methyltransferase family members and the complex regulatory role of methylation in the nervous system.

In *Drosophila, TRMT9B* transcription peaks in late embryogenesis shortly before synaptogenesis begins in the developing nervous system, and TRMT9B is primarily expressed in the nervous system, with lower levels of expression in carcass (muscle and epidermis), fat body, imaginal discs, and testes (Brown et al., 2014). Similarly, human *TRMT9B* is highly expressed in the nervous system, including the cerebellum, visual cortex, hippocampus and cerebral cortex (Uhlén et al., 2015). Nonetheless, to date TRMT9B has only been studied in the context of non-neuronal cancers, although a recent study found that TRMT9B expression is significantly increased along with a host of gene regulatory proteins during NeuroD1-mediated conversion of astrocytes to neurons (Ma et al., 2022). Another RNAi screen in *Drosophila* identified a putative role for *Drosophila TRMT9B* in promoting dendrite outgrowth in class IV multidendritic sensory neurons and thermal nociception (Honjo et al., 2016). We find TRMT9B negatively regulates synapse formation at the NMJ and, despite synaptic overgrowth, *TRMT9B* mutants display a significant impairment in neurotransmitter release. This suggests that abnormal synaptic transmission in *TRMT9B* mutants is due to functional impairment of the individual synapses that compose the NMJ rather than altered synapse number, pointing to a broad requirement for methylation of TRMT9B substrates in nervous system development.

In 2004, TRMT9B was first identified as a potential tumor suppressor in colorectal cancer (Flanagan et al., 2004). Since that time, reduced TRMT9B expression due to genomic rearrangements or epigenetic silencing has been observed in ovarian, lung, and other carcinomas (Begley et al., 2013; Chen et al., 2017; Flanagan et al., 2004; Wang et al., 2018). Restoration of TRMT9B expression in colon, lung, and ovarian cancer cells reduces proliferation and significantly reduces colon tumor growth *in vivo*, confirming an important tumor suppressor role for TRMT9B (Begley et al., 2013; Chen et al., 2017; Wang et al., 2018). Despite the numerous links to cancer, the normal biological role of TRMT9B function has remained unknown. Although insight into TRMT9B’s biochemical role has been limited by the lack of an animal model, a gain-of-function study found that TRMT9B promotes expression of tumor suppressor LIN9, a core member of the DREAM complex that represses cell cycle-dependent gene expression, and blocks HIF1-α-dependent adaptation to a hypoxia (Begley et al., 2013). A more recent study found that stress-dependent phosphorylation modulates TRMT9B’s interactions with the 14-3-3 gamma, epsilon, and eta signaling molecules and its ability to suppress proliferation and hypoxic adaptation (Gu et al., 2018).

The tRNA methyltransferase (Trm) family of enzymes has expanded from 18 Trm proteins that form 15 holoenzymes in yeast to 34 Trm homologs in animals, with all yeast tRNA methyltransferases represented in animals and more than half represented by two or more paralogs in humans (Towns and Begley, 2012). Many of the additional family members in animals are yet to be characterized, raising the question of whether they maintain redundant roles as canonical tRNA methyltransferases or have evolved new functions. Of the two metazoan homologs of yeast Trm9, mammalian ALKBH8 has been shown to methylate tRNA wobble uridines (Fu et al., 2010a; Kalhor and Clarke, 2003; Songe-Moller et al., 2010). Our quantitative mass spectrometry studies demonstrate that ALKBH8 is similarly responsible for the methylation of tRNA wobble uridines in *Drosophila*. Interestingly, ALKBH8 has recently been linked to intellectual disability in several families, and, we have found, acts through a tRNA wobble uridine-dependent mechanism to regulate nervous system development (Maddirevula et al., 2022; Monies et al., 2019; Saad et al., 2021; Madhawandi et al., in preparation).

In contrast, we find that TRMT9B is either dispensable or plays a conditional role in tRNA wobble uridine methylation not detected in our analysis. Our findings are consistent with the previous observation that ALKBH8, but not TRMT9B, can rescue tRNA methylation in a yeast deletion of Trm9 (Begley et al., 2013) and raise the possibility that TRMT9B may have evolved a new function. TRMT9B might also function as a tRNA wobble uridine methyltransferase under specific conditions or methylate only one or a few, possibly brain-enriched, tRNAs as has recently been demonstrated for a mammalian homolog of yeast Trm140, which methylates tRNAs in the anticodon loop (Xu et al., 2017). This possibility is supported by the recent finding that overexpression of TRMT9B leads to subtle, but significant changes in tRNAs with an mcm5U terminal modification, although mcm5S2U levels remain unchanged (Jungfleisch et al., 2022).

To explore how TRMT9B might function biochemically, we investigated TRMT9B’s sequence and structural homology with SAM-dependent methyltransferases. Several lines of evidence suggest that TRMT9B functions as a methyltransferase: (1) residues required for enzymatic function in yeast are highly conserved, including the canonical Class I SAM-dependent methyltransferase GXGXGX motif and acidic amino acid at the end of the second beta strand, both of which mediate SAM binding; (2) structural modeling reveals remarkable similarity between the methyltransferase domains of yeast Trm9 and fly and human TRMT9B and ALKBH8; and (3) disruption of two residues critical for methyltransferase function eliminates the ability of a TRMT9B transgene to rescue synaptic growth without disrupting its ability to fold or localize properly. Together, these findings support the conclusion that TRMT9B regulates nervous system development through a methyltransferase-dependent mechanism.

The next major step will be identifying TRMT9B’s substrate(s). A previous study identified changes to a number of tRNA modifications upon expression of TRMT9B (Begley et al., 2013). However, it is unclear if any of these modifications are direct targets of TRMT9B as many tRNA modifications depend on prior modifications. To investigate the possibility that TRMT9B catalyzes a distinct modification on tRNAs, we assessed a panel of tRNA post-transcriptional modifications in our mutants via quantitative mass spectrometry. We did not identify a TRMT9B-dependent modification among our panel, which includes nine of the 12 modifications identified by Begley et al., 2013. However, it remains possible that TRMT9B catalyzes a modification we were unable to evaluate, including an animal-specific tRNA modification yet to be identified. As most of our understanding of tRNA methylation comes from studies in single-cell organisms, it would not be surprising if additional tRNA modifications specific to animals, and possibly enriched in specific tissues such as the nervous system, remain undiscovered. Another possibility is that TRMT9B methylates a non-tRNA substrate. In fact, several members of the expanded metazoan tRNA methyltransferase family methylate novel substrates. For example, the NSUN family of yeast Trm4 homologs, methylate not only tRNAs, but mRNA, rRNAs and noncoding RNAs, and have important roles in both cancer and neurodevelopment (Abbasi-Moheb et al., 2012; Chellamuthu and Gray, 2020; Chen et al., 2021). Intriguingly, we have found that both fly and human TRMT9B immunoprecipitate TRMT112, the homolog of obligate yeast Trm9 co-factor Trm112, in IP-LC-MS/MS experiments (2 unique peptides, 11% peptide coverage; (Gu et al., 2018), respectively). TRMT112 promotes the methylation of diverse substrates as an obligate cofactor for several tRNA, rRNA, and protein methyltransferases – an indication of the expansive landscape of potential TRMT9B substrates (Gao et al., 2020; Garcia et al., 2021; Guy and Phizicky, 2014; van Tran et al., 2019; Yang et al., 2021). A recent study of the Trm112 interactome in archea identified a similarly broad array of interacting methyltransferases as well as small molecule methyltransferases involved in steroid and vitamin biosynthesis (van Tran et al., 2018), raising still more diverse possibilities. Especially exciting is the possibility hinted at by its endogenous expression pattern that TRMT9B methylates specific targets in different cellular and sub-cellular contexts, including synaptic targets. Given TRMT9B’s fundamental roles as a regulator of synapse formation and function and a tumor suppressor, it will be of great interest and clinical significance to identify target(s) of TRMT9B methylation in future studies.

## Materials and methods

### Drosophila genetics and gene editing

The following stocks used in this study are available through the Bloomington Drosophila Stock Center (BDSC): *w*^*1118*^ (RRID:BDSC_5905), *vasa-Cas9* (RRID:BDSC_51324), piggyBac transposase (RRID:BDSC_8283), *attP2* (RRID:BDSC_25710), *elav*^*c155*^ *Gal4* (RRID:BDSC_458), *24B Gal4* (RRID:BDSC_92172). y^1^ w^*^; Mi{MIC}fid^MI13909^ (*TRMT9B MiMIC*^*Stop*^) and *y*^*1*^ *w*^*\**^; *Mi{PT-GFSTF*.*1}dfid*^*MI13909-GFSTF*.*1*^ (*TRMT9B MiMIC*^*GFP*^) were generated through the Gene Disruption Project and obtained from the BDSC (RRID:BDSC_59229 and RRID:BDSC_66771) (Nagarkar-Jaiswal et al., 2015; Venken et al., 2011). A lab *w*^*1118*^ strain that is the genetic background of the CRISPR mutants was used as a control unless otherwise noted. Full genotypes for all lines used in this study are listed in Tables S1 and S2.

CRISPR-based homology directed repair strategies were used to generate knock-outs and in-frame endogenous tags. Target sites were selected using CRISPR Optimal Target Finder program (http://targetfinder.flycrispr.neuro.brown.edu; Gratz et al., 2014). The following gRNAs were used: TRMT9B^KO^ 5′-AATCCATCGTCCTGAAATAG-3′, 5’-AGTCTATTACTCTAGTCGGC-3’ TRMT9B^HA+IC^ 5′-CGACGGCTTTGTTGAATGCG-3′. gRNA and donor plasmids were generated as described in (Gratz et al., 2014). *Vasa-Cas9* embryos were injected with a mixture of two gRNA plasmids (100ng/μL, each) and a double stranded DNA donor plasmid (500ng/μL) by BestGene, Inc., crossed to *w*^*1118*^ flies after eclosion, and progeny screened for DsRed expression in the eye. Knock-out alleles were generated by replacing endogenous loci with a visible marker and attP landing site (Gratz et al., 2013). Endogenously tagged alleles were created using a scarless CRISPR-piggyBac approach (https://flycrispr.org; Bruckner et al., 2017). An HA or superfolder GFP (sfGFP) tag flanked by flexible linkers and a visible marker flanked by piggyBac inverted terminal repeat sequences were inserted immediately downstream of the sole *TRMT9B* or ALKBH8 (CG17807) translational start sites. This generates null alleles due stop codons in the inverted repeat sequences. TRMT9B lines were crossed to piggyBac transposase to remove the marker cassette and generate in-frame tags. All engineered lines were confirmed by PCR and Sanger sequencing.

UAS rescue lines were generated by cloning full-length *Drosophila* and human TRMT9B coding sequence into pUAST-C5 (*Drosophila* Genomics Resource Center #1261). To generate methyltransferase-dead *Drosophila* TRMT9B, mutations to critical residues in the methyltransferase domain were introduced via KLD reaction. Specifically, we mutated codon 417 (GAG to GCC) and codon 693 (CAC to GCC) to introduce E139A and H231A mutations. These amino acids correspond to yeast residues D72 and H116, which were identified as critical for methyltransferase function (Létoquart et al., 2015). In parallel, we generated V5 tagged wild-type and methyltransferase-dead *Drosophila* TRMT9B transgenes to confirm localization. All transgenes were integrated into the attP2 landing site by BestGene, Inc. (Groth et al., 2004).

### Immunostaining and confocal imaging

Male third instar larvae were dissected in Ca^2+^-free saline and fixed for 10 minutes with 4% paraformaldehyde in PBS. Dissected larvae were washed and permeabilized in PBS with 0.1% Triton-X then blocked overnight at 4°C in PBS containing 0.1% Triton-X and 1% BSA. Dissected larvae were incubated with primary antibodies for two hours at room temperature and secondary antibodies for one hour at room temperature, then mounted in Vectashield (Vector Laboratories). For localization studies, male third-instar larvae were dissected in ice-cold Ca^2+^-free saline and fixed for 6 min in Bouin’s Fixative. Dissected larvae were washed and permeabilized in PBS with 0.1% Triton-X then blocked overnight at 4°C in PBS containing 0.1% Triton-X, 5%NGS and 1% BSA, followed by overnight incubation with primary and secondary antibodies, then mounted in Vectashield (Vector Laboratories). The following antibodies were used at the indicated concentrations: mouse anti-Bruchpilot (Brp) at 1:100 (Developmental Studies Hybridoma Bank (DSHB) #nc82; RRID: AB_2314866), rabbit anti-HA at 1:500 (Cell Signaling Technology C29F4), and anti-HRP conjugated to Alexa Fluor 647 at 1:500 (Jackson ImmunoResearch Laboratories, Inc., #123-605-021; RRID:AB_2338967), rabbit anti-GFP conjugated to AlexaFluor 488 at 1:500 (Thermo Fisher Scientific, #A-21311; RRID:AB_221477), and secondary antibodies Alexa Fluor 488 and Alexa Fluor 568 at 1:500 (Thermo Fisher Scientific). Images were obtained on a Nikon A1R HD confocal microscope with Plan Apo 40x and 60X 1 oil-immersion objectives at a pixel resolution of 130 nm.

### Electrophysiology

Electrophysiological baseline readings were performed on male third instar larvae. Larvae were dissected in low Ca^2+^ modified hemolymph-like saline (HL3 in mM: 70 NaCl, 5 KCl, 20 MgCl2, 10 NaHCO3, 115 sucrose, 5 trehalose, 5 HEPES and 0.2 Ca^2+^, pH 7.2). Excitatory junction potentials were recorded using 1.0MM x 0.58MM, 4” borosilicate glass sharp electrode (electrode resistance between 10 and 15 MΩ) filled with 3M KCl. Recordings were performed in hemolymph-like saline solution containing 0.6 mM Ca^2+^ on muscle 6 of abdominal segments A3 and A4. A 1.5MM x 1.12MM, 4” borosilicate polished glass electrode was used to suction nerves. Recordings were performed on a Nikon FN1 microscope using a 40x (0.80 NA) water-dipping objective, and acquired using an AxoClamp 900A amplifier, Axon Digidata 1550B low-noise data acquisition system, and pClamp 11.0.3 software (Molecular Devices). Electrophysiological sweeps were digitized at 10 kHz, and filtered at 0.1 kHz. Miniature excitatory junctional potentials (mEJPs) were recorded with no external stimulation. For each recording, at least 100 mEJPs were analyzed using Mini Analysis (Synaptosoft) to obtain a mean mEJP amplitude value per muscle. Cut motor axons of each segment were stimulated for 0.5ms with an A-M Systems Isolated pulse stimulator to elicit EJPs. Stimulus intensity was adjusted to consistently elicit compound responses from both type Ib and Is motorneurons. At least 30 consecutive EJPs were recorded for each cell and analyzed in pClamp to obtain mean amplitude. Quantal content (QC) was determined for each recording by calculating the ratio of mean EJP amplitude to mean mEJP amplitude then averaging recordings across all NMJs for a given genotype. Muscle input resistance (Rin) and resting membrane potential (V_rest_) were monitored during each experiment. Recordings were only performed if the V_rest_ was between -55 and -80 mV and Rin was 5 MΩ or higher.

### Sequence alignment, phylogeny, and homology modeling

Alignments of the methyltransferase domains as defined by Pfam (Mistry et al., 2021) were conducted using T-Coffee (Notredame et al., 2000) using the following UniProt sequences: P49957 (*Saccharomyces cerevisiae* Trm9), Q9VBJ3 (*Drosophila* TRMT9B), Q9P272 (human TRMT9B), Q9W232 (*Drosophila* ALKBH8), and Q96BT7 (human ALKBH8). A phylogenetic tree was determined by PhyML maximum likelihood and visualized with TreeDyn in phylogeny.fr (Dereeper et al., 2008; Lemoine et al., 2019). Template-based structural homology models of the methyltransferase domains of *Drosophila* and human TRMT9B and ALKBH8 were performed using Modeller (Pieper et al., 2014) through the ModWeb online service (https://modbase.compbio.ucsf.edu/modweb). Best scoring models for all four proteins included *Yarrowia lipolytica* Trm9, obtained from 2.5 Å X-ray diffraction of Trm9-Trm112, PDB: 5CM2 chain Z (Létoquart et al., 2015). All homology models built with PDB 5CM2 chain Z were reliable with GA341 score of >0.99989 and normalized DOPE (zDOPE) score < 0). *Drosophila* and human TRMT9B models were built with residues 39 to 160 of *Y. lipolytica*, with 39% and 46% sequence identity, respectively. The *Drosophila* ALKBH8 model was built with residues 39 to 191 with 40% sequence identity and the human ALKBH8 model was built with residues 16 to 174 with 46% sequence identity. To model the mutations in methyltransferase dead TRMT9B, we again used residues 39 to 160. Additional details on the best scoring models can be found in the protein data bank files (Files S1-S5). Models were visualized and compared to PDB 5CM2 chain Z using UCSF ChimeraX (Pettersen et al., 2021). A template-free model for *Drosophila* TRMT9B was obtained from AlphaFold version 1 predictions and is provided in the supplemental files (File S6, Jumper et al., 2021; Varadi et al., 2022). The AlphaFold model confidently predicted (pLDDT > 70) the methyltransferase domain, residues 123-278, although a biological function was not defined. Confidence scores for critical methyltransferase activity including the GxGxGx domain, the acidic residue, and SAM binding histidine were very high (pLDDT between 89.8 - 95.7).

### Liquid chromatography-mass spectrometry of RNA modifications

Vials containing adult fly heads were flash frozen in liquid nitrogen and stored at −80°C. Twenty-five milligrams of frozen adult flies were homogenized with a plastic pestle in a 1.5 mL microfuge tube while adding 250 μL of TRIzol reagent at a time until 1 mL was added. RNA was then extracted via an adapted TRIzol extraction protocol (Bogart and Andrews, 2006). RNA was resuspended in RNase-free ddH_2_O. Total RNA was processed and analyzed by LC-MS as previously described (Su et al., 2014; Zhang et al., 2020). Briefly, total RNA (20 ug) was digested and dephosphorylated with Benzonase, Phosphodiesterase, and Calf Intestinal phosphatase for 3 hours at 37°C. Enzymes and undigested RNAs were removed using a 10,000-Da MWCO spin filter (Amicon) and purified ribonucleosides were stored at -80°C until further analysis. Ribonucleosides were separated using a Hypersil GOLD™ C18 Selectivity Column (Thermo Scientific) followed by nucleoside analysis using a Q Exactive Plus Hybrid Quadrupole-Orbitrap. The modification difference ratio was calculated using the m/z intensity values of each modified nucleoside between control and mutants following normalization to the sum of intensity values for the canonical nucleosides A, U, G and C.

### IP-MS analysis of TRMT9B-interacting proteins

We pulled down endogenous TRMT9B protein and interacting proteins in *y*^*1*^ *w*^*\**^; *Mi{PT-GFSTF*.*1}dfid*^*MI13909-GFSTF*.*1*^, in which TRMT9B is tagged with GFP and FLAG (Nagarkar-Jaiswal et al., 2015). Whole adult flies were flash frozen in liquid nitrogen and manually homogenized. Cells were then lysed in lysis buffer (50 mM Tris-HCl, pH 7.5, 150 mM NaCl, 1 mM EDTA, 1% Triton X-100 supplemented with a protease inhibitor cocktail). Lysates were cleared by centrifugation and immunoprecipitated with anti-FLAG-antibody-conjugated agarose beads. Beads were washed three times with FLAG-wash buffer (50mM Tris-HCl, pH 7.5, 500mM NaCl, 1mM EDTA, 1% Triton X-100) and twice with TBS buffer (50 mM Tris-HCl, pH 7.5, 150 mM NaCl). EGFP-FIAsH-StrepII-3xFLAG - tagged TRMT9B was eluted with 100 μg/ml of 3×FLAG peptide in TBS buffer. As a control, an identical purification was performed in w1118 control flies. A silver stain was performed to confirm the presence of proteins in the eluted fractions, which were then subjected to trypsin digestion and tandem mass spectrometry. We used a 1% false discovery rate peptide threshold and 99% protein threshold to identify interacting proteins identified by at least two unique peptides.

### Experimental Design and Statistical Analysis

Quantifications were conducted blind to genotype. Statistical analyses were conducted in GraphPad Prism 8 and 9. ANOVA followed by Tukey’s test was used for multiple comparisons of normally distributed data. Multiple comparisons of non-normally distributed data were performed using the Kruskal-Wallis test followed by Dunn’s multiple comparisons test. Significance is reported as values less than 0.05, 0.01, 0.001, and 0.ooo1 represented by one, two, three, or four stars, respectively.Unless otherwise noted, significance refers to the indicated genotype compared to control. Error bars represent the mean with SEM. Full genotypes, sample number, p values, and absolute values for bouton quantifications and electrophysiology are presented in Tables S1 and S2, respectively.

## Acknowledgements

This work was supported by a grant from the NIH National Institute of Neurological Disorders and Stroke to K.M.O.-G. and D.F. (R01 NS117068) and trainee support from the National Institute of General Medical Sciences to C.A. Hogan and K.R. Madhwani through the University of Wisconsin Predoctoral Training Program in Genetics (T32GM007133) and J.L. Dumouchel through the Brown University Predoctoral Training Program in Trans-disciplinary Pharmacological Sciences (T32 GM077995). We thank Sina Ghaemmaghami and the Mass Spectrometry Resource Lab at the University of Rochester; the Biotechnology Center Mass Spectrometry/Proteomics Facility at the University of Wisconsin-Madison; the Laboratories of Genetics and Cell and Molecular Biology at the University of Wisconsin-Madison where a subset of this work was completed; and the Developmental Studies Hybridoma Bank and Bloomington Drosophila Stock Center for providing antibodies and fly stocks. We thank Sebastien Santini (CNRS/AMU IGS UMR7256) and the PACA Bioinfo platform (supported by IBISA) for the availability and management of the phylogeny.fr website used for phylogeny and sequence alignment. Molecular graphics images were created using the UCSF Chimera package from the Resource for Biocomputing, Visualization, and Informatics at the University of California, San Francisco (supported by NIH P41 RR001081). We are grateful to the O’Connor-Giles and Fu labs and Heather Broihier for critical feedback on the work and manuscript.

---

**Table S1.** Absolute values and statistics for synaptic growth data. Full genotypes, raw bouton counts ± SEM and the number of NMJs quantified are provided for each panel of Figures 1, 2, 5, and S1. P values and significance represent comparisons to control unless noted.

| Figure | Genotype | Bouton # | n (NMJs) | P value (significance) |
| --- | --- | --- | --- | --- |
| 1A | *W<sup>1118</sup>* | 29.0 ± .89 | 44 | |
| 1B | *TRMT9B<sup>KO</sup>* | 39.2 ± 1.2 | 40 | <0.0001 (****) |
| 1C | *TRMT9B<sup>HA+IC</sup>* | 39.1 ± 1.0 | 39 | <0.0001 (****)<br>*TRMT9B<sup>HA+IC</sup>* vs *TRMT9B<sup>HA</sup>*<br>0.0038 (**) |
| 1D | *TRMT9B<sup>KO/HA+IC</sup>* | 38.0 ± .91 | 42 | <0.0001 (****) |
| 1E | *TRMT9B<sup>HA</sup>* | 33.6 ± 1.0 | 39 | 0.0335 (*) |
| 2A | *C155-Gal4/+* | 32.5 ± 1.5 | 26 | |
| 2B | *TRMT9B<sup>KO/HA+IC</sup>* | 36.9 ± 1.2 | 36 | 0.0240 (*) |
| 2C | *C155-Gal4; TRMT9B<sup>KO</sup>/UAS-TRMT9B, TRMT9B<sup>HA+IC</sup>* (C155 Rescue) | 41.6 ± 1.8 | 30 | <0.0001 (****)<br>C155 Rescue vs *TRMT9B<sup>KO/HA+IC</sup>*<br>0.1049 (ns) |
| 2E | *24B-Gal4/+* | 29.5 ± 1.0 | 41 | |
| 2F | *TRMT9B<sup>KO/HA+IC</sup>* | 37.7 ± 0.8 | 42 | <0.0001 (****) |
| 2G | *24B-Gal4, TRMT9B<sup>KO</sup>/UAS-TRMT9B, TRMT9B<sup>HA+IC</sup>* (24B Rescue) | 31.98 ± 0.8 | 40 | 0.6437 (ns)<br>24B Rescue vs *TRMT9B<sup>KO/HA+IC</sup>*<br>0.0004 (****) |
| 5A | *24B-Gal4/+* | 31.4 ± 0.7 | 60 | |
| 5B | *TRMT9B<sup>KO/HA+IC</sup>* | 35.4 ± 0.7 | 59 | 0.0003 (***)<br>*TRMT9B<sup>KO/HA+IC</sup>* vs 24B *TRMT9B* Rescue <0.0001 (****) |
|  |  |  |  | *TRMT9B*<sup>KO/HA+IC</sup> vs 24B<br>*TRMT9B*<sup>MTD</sup> Rescue 0.5496 (ns) |
| 5C | 24B-Gal4, *TRMT9B*<sup>KO</sup>/UAS-*TRMT9B*, *TRMT9B*<sup>HA+IC</sup> (24B *TRMT9B* Rescue) | 30.3 ± 0.8 | 57 | 0.7030 (ns)<br>24B *TRMT9B* Rescue vs 24B *TRMT9B*<sup>MTD</sup> Rescue 0.0010 (***) |
| 5D | 24B-Gal4, *TRMT9B*<sup>KO</sup>/UAS-*TRMT9B*<sup>MTD</sup>, *TRMT9B*<sup>HA+IC</sup> (24B *TRMT9B*<sup>MTD</sup> Rescue) | 34.1 ± 0.7 | 59 | 0.0296 (*) |
| S1A | Control | 27.5 ± .74 | 46 |  |
| S1A | *TRMT9B*<sup>KO</sup> | 38.0 ± 1.4 | 66 | <0.0001 (****) |
| S1A | *TRMT9B*<sup>KO</sup>/Df | 40.2 ± 1.0 | 53 | <0.0001 (****) |
| S1A | *TRMT9B* MiMIC<sup>stop</sup> | 39.3 ± 1.1 | 58 | <0.0001 (****)<br>*TRMT9B* MiMIC<sup>stop</sup> vs *TRMT9B* MiMIC<sup>[GFP/Stop]</sup> 0.0005 (***) |
| S1A | *TRMT9B* MiMIC<sup>[GFP/Stop]</sup> | 32.9 ± .74 | 49 | 0.0037 (**) |

**Table S2.**
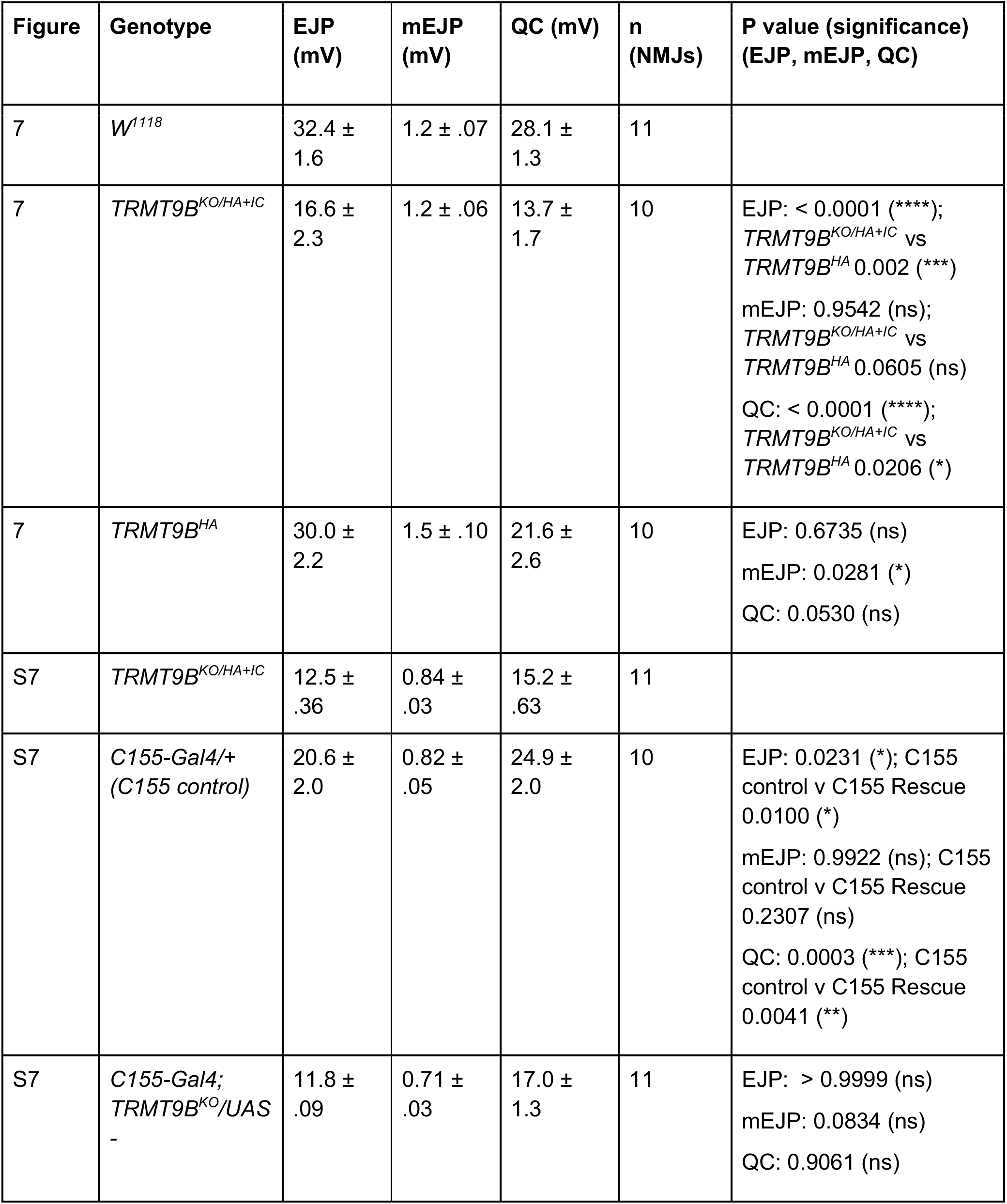
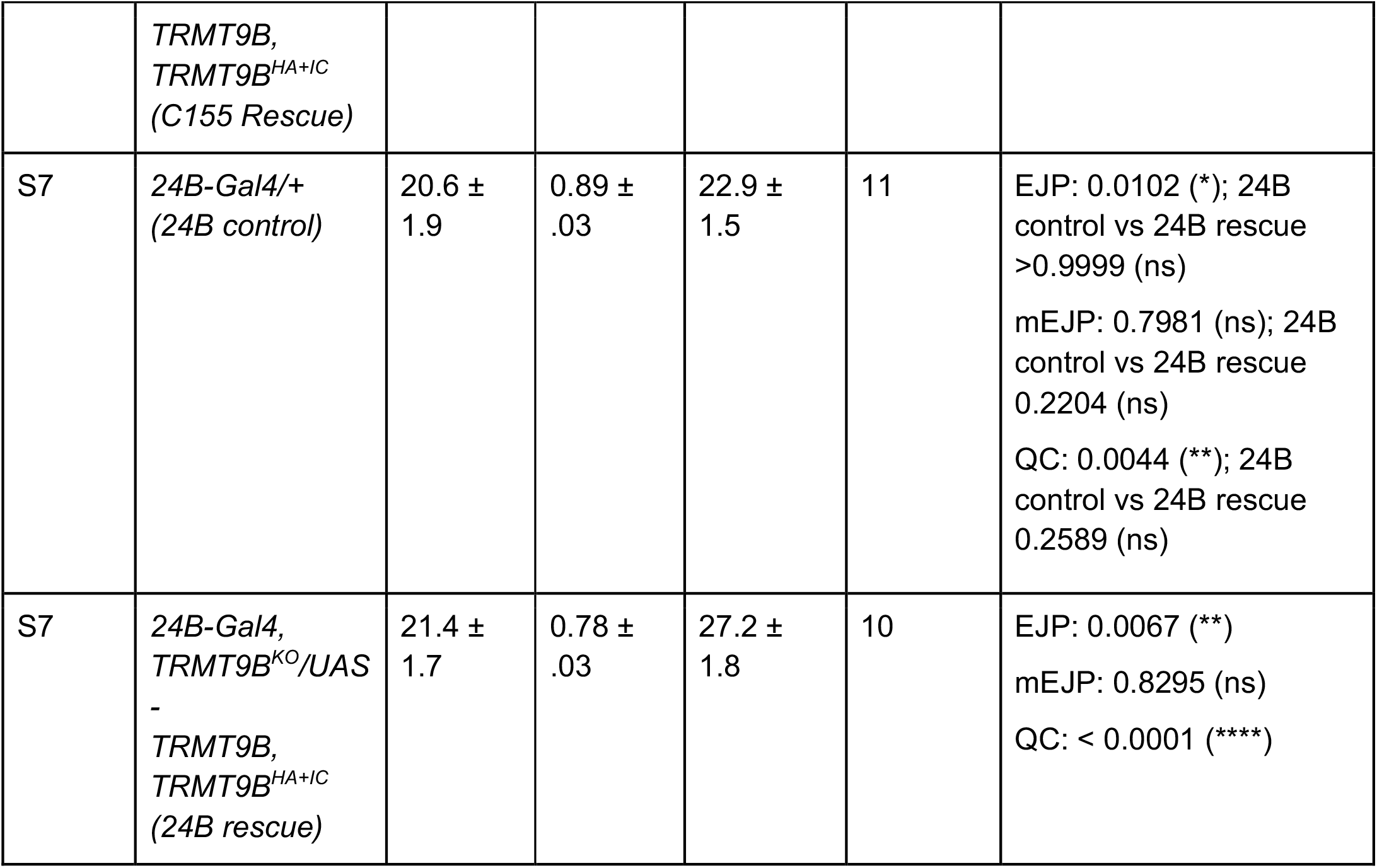
Absolute values and statistics for electrophysiological data. Full genotypes and mEJP, EJP and QC ± SEM are provided along with the number of NMJs recorded. P values and significance represent comparisons to control unless noted.

## References

Abbasi-Moheb, L., S. Mertel, M. Gonsior, L. Nouri-Vahid, K. Kahrizi, S. Cirak, D. Wieczorek, M.M. Motazacker, S. Esmaeeli-Nieh, K. Cremer, R. Weißmann, A. Tzschach, M. Garshasbi, S.S. Abedini, H. Najmabadi, H.H. Ropers, S.J. Sigrist, and A.W. Kuss. 2012. Mutations in NSUN2 cause autosomal-recessive intellectual disability. Am J Hum Genet. 90:847–855.

Agris, P.F., A. Narendran, K. Sarachan, V.Y.P. Väre, and E. Eruysal. 2017. The Importance of Being Modified: The Role of RNA Modifications in Translational Fidelity. Enzymes. 41:1–50.

Begley, U., M. Dyavaiah, A. Patil, J.P. Rooney, D. DiRenzo, C.M. Young, D.S. Conklin, R.S. Zitomer, and T.J. Begley. 2007. Trm9-catalyzed tRNA modifications link translation to the DNA damage response. Molecular cell. 28:860–870.

Begley, U., M.S. Sosa, A. Avivar-Valderas, A. Patil, L. Endres, Y. Estrada, C.T. Chan, D. Su, P.C. Dedon, J.A. Aguirre-Ghiso, and T. Begley. 2013. A human tRNA methyltransferase 9-like protein prevents tumour growth by regulating LIN9 and HIF1-alpha. EMBO molecular medicine. 5:366–383.

Berman, H.M., J. Westbrook, Z. Feng, G. Gilliland, T.N. Bhat, H. Weissig, I.N. Shindyalov, and P.E. Bourne. 2000. The Protein Data Bank. Nucleic acids research. 28:235–242.

Boccaletto, P., F. Stefaniak, A. Ray, A. Cappannini, S. Mukherjee, E. Purta, M. Kurkowska, N. Shirvanizadeh, E. Destefanis, P. Groza, G. Avşar, A. Romitelli, P. Pir, E. Dassi, S.G. Conticello, F. Aguilo, and J.M. Bujnicki. 2022. MODOMICS: a database of RNA modification pathways. 2021 update. Nucleic acids research. 50:D231–d235.

Bogart, K., and J. Andrews 2006. Extraction of Total RNA from Drosophila. CGB Technical Report. 2006–10.

Brown, J.B., N. Boley, R. Eisman, G.E. May, M.H. Stoiber, M.O. Duff, B.W. Booth, J. Wen, S. Park, A.M. Suzuki, K.H. Wan, C. Yu, D. Zhang, J.W. Carlson, L. Cherbas, B.D. Eads, D. Miller, K. Mockaitis, J. Roberts, C.A. Davis, E. Frise, A.S. Hammonds, S. Olson, S. Shenker, D. Sturgill, A.A. Samsonova, R. Weiszmann, G. Robinson, J. Hernandez, J. Andrews, P.J. Bickel, P. Carninci, P. Cherbas, T.R. Gingeras, R.A. Hoskins, T.C. Kaufman, E.C. Lai, B. Oliver, N. Perrimon, B.R. Graveley, and S.E. Celniker. 2014. Diversity and dynamics of the Drosophila transcriptome. Nature. 512:393–399.

Bruckner, J.J., H. Zhan, S.J. Gratz, M. Rao, F. Ukken, G. Zilberg, and K.M. O’Connor-Giles. 2017. Fife organizes synaptic vesicles and calcium channels for high-probability neurotransmitter release. The Journal of cell biology. 216:231–246.

Cai, W.M., Y.H. Chionh, F. Hia, C. Gu, S. Kellner, M.E. McBee, C.S. Ng, Y.L. Pang, E.G. Prestwich, K.S. Lim, I.R. Babu, T.J. Begley, and P.C. Dedon. 2015. A Platform for Discovery and Quantification of Modified Ribonucleosides in RNA: Application to Stress-Induced Reprogramming of tRNA Modifications. Methods Enzymol. 560:29–71.

Chellamuthu, A., and S.G. Gray. 2020. The RNA Methyltransferase NSUN2 and Its Potential Roles in Cancer. Cells. 9.

Chen, C., B. Huang, J.T. Anderson, and A.S. Byström. 2011. Unexpected accumulation of ncm(5)U and ncm(5)S(2) (U) in a trm9 mutant suggests an additional step in the synthesis of mcm(5)U and mcm(5)S(2)U. PLoS One. 6:e20783.

Chen, H.M., J. Wang, Y.F. Zhang, and Y.H. Gao. 2017. Ovarian cancer proliferation and apoptosis are regulated by human transfer RNA methyltransferase 9-likevia LIN9. Oncology letters. 14:4461–4466.

Chen, Y.S., W.L. Yang, Y.L. Zhao, and Y.G. Yang. 2021. Dynamic transcriptomic m(5) C and its regulatory role in RNA processing. Wiley Interdiscip Rev RNA. 12:e1639.

Citri, A., and R.C. Malenka. 2008. Synaptic plasticity: multiple forms, functions, and mechanisms. Neuropsychopharmacology. 33:18–41.

GTEx Consortium. 2015. Human genomics. The Genotype-Tissue Expression (GTEx) pilot analysis: multitissue gene regulation in humans. Science. 348:648–660.

Dereeper, A., V. Guignon, G. Blanc, S. Audic, S. Buffet, F. Chevenet, J.F. Dufayard, S. Guindon, V. Lefort, M. Lescot, J.M. Claverie, and O. Gascuel. 2008. Phylogeny.fr: robust phylogenetic analysis for the non-specialist. Nucleic acids research. 36:W465–469.

Dewe, J.M., B.L. Fuller, J.M. Lentini, S.M. Kellner, and D. Fu. 2017. TRMT1-Catalyzed tRNA Modifications Are Required for Redox Homeostasis To Ensure Proper Cellular Proliferation and Oxidative Stress Survival. Molecular and cellular biology. 37.

Dietzl, G., D. Chen, F. Schnorrer, K.C. Su, Y. Barinova, M. Fellner, B. Gasser, K. Kinsey, S. Oppel, S. Scheiblauer, A. Couto, V. Marra, K. Keleman, and B.J. Dickson. 2007. A genome-wide transgenic RNAi library for conditional gene inactivation in Drosophila. Nature. 448:151–156.

Flanagan, J.M., S. Healey, J. Young, V. Whitehall, D.A. Trott, R.F. Newbold, and G. Chenevix-Trench. 2004. Mapping of a candidate colorectal cancer tumor-suppressor gene to a 900-kilobase region on the short arm of chromosome 8. Genes Chromosomes Cancer. 40:247–260.

Fu, D., J.A. Brophy, C.T. Chan, K.A. Atmore, U. Begley, R.S. Paules, P.C. Dedon, T.J. Begley, and L.D. Samson. 2010a. Human AlkB homolog ABH8 Is a tRNA methyltransferase required for wobble uridine modification and DNA damage survival. Molecular and cellular biology. 30:2449–2459.

Fu, Y., Q. Dai, W. Zhang, J. Ren, T. Pan, and C. He. 2010b. The AlkB domain of mammalian ABH8 catalyzes hydroxylation of 5-methoxycarbonylmethyluridine at the wobble position of tRNA. Angewandte Chemie. 49:8885–8888.

Gao, J., B. Wang, H. Yu, G. Wu, C. Wan, W. Liu, S. Liao, L. Cheng, and Z. Zhu. 2020. Structural insight into HEMK2-TRMT112-mediated glutamine methylation. Biochem J. 477:3833–3838.

Garcia, B.C.B., M. Horie, S. Kojima, A. Makino, and K. Tomonaga. 2021. BUD23-TRMT112 interacts with the L protein of Borna disease virus and mediates the chromosomal tethering of viral ribonucleoproteins. Microbiol Immunol. 65:492–504.

Gratz, S.J., A.M. Cummings, J.N. Nguyen, D.C. Hamm, L.K. Donohue, M.M. Harrison, J. Wildonger, and K.M. O’Connor-Giles. 2013. Genome engineering of Drosophila with the CRISPR RNA-guided Cas9 nuclease. Genetics. 194:1029–1035.

Gratz, S.J., F.P. Ukken, C.D. Rubinstein, G. Thiede, L.K. Donohue, A.M. Cummings, and K.M. O’Connor-Giles. 2014. Highly specific and efficient CRISPR/Cas9-catalyzed homology-directed repair in Drosophila. Genetics. 196:961–971.

Graveley, B.R., A.N. Brooks, J.W. Carlson, M.O. Duff, J.M. Landolin, L. Yang, C.G. Artieri, M.J. van Baren, N. Boley, B.W. Booth, J.B. Brown, L. Cherbas, C.A. Davis, A. Dobin, R. Li, W. Lin, J.H. Malone, N.R. Mattiuzzo, D. Miller, D. Sturgill, B.B. Tuch, C. Zaleski, D. Zhang, M. Blanchette, S. Dudoit, B. Eads, R.E. Green, A. Hammonds, L. Jiang, P. Kapranov, L. Langton, N. Perrimon, J.E. Sandler, K.H. Wan, A. Willingham, Y. Zhang, Y. Zou, J. Andrews, P.J. Bickel, S.E. Brenner, M.R. Brent, P. Cherbas, T.R. Gingeras, R.A. Hoskins, T.C. Kaufman, B. Oliver, and S.E. Celniker. 2011. The developmental transcriptome of Drosophila melanogaster. Nature. 471:473–479.

Greenberg, M.V.C., and D. Bourc’his. 2019. The diverse roles of DNA methylation in mammalian development and disease. Nat Rev Mol Cell Biol. 20:590–607.

Groth, A.C., M. Fish, R. Nusse, and M.P. Calos. 2004. Construction of transgenic Drosophila by using the site-specific integrase from phage phiC31. Genetics. 166:1775–1782.

Gu, C., J. Ramos, U. Begley, P.C. Dedon, D. Fu, and T.J. Begley. 2018. Phosphorylation of human TRM9L integrates multiple stress-signaling pathways for tumor growth suppression. Science advances. 4:eaas9184.

Guang, S., N. Pang, X. Deng, L. Yang, F. He, L. Wu, C. Chen, F. Yin, and J. Peng. 2018. Synaptopathology Involved in Autism Spectrum Disorder. Frontiers in cellular neuroscience. 12:470.

Guy, M.P., and E.M. Phizicky. 2014. Two-subunit enzymes involved in eukaryotic post-transcriptional tRNA modification. RNA biology. 11:1608–1618.

Honjo, K., S.E. Mauthner, Y. Wang, J.H. Skene, and W.D. Tracey, Jr. 2016. Nociceptor-Enriched Genes Required for Normal Thermal Nociception. Cell reports. 16:295–303.

Hu, Y., A. Comjean, C. Roesel, A. Vinayagam, I. Flockhart, J. Zirin, L. Perkins, N. Perrimon, and S.E. Mohr. 2017. FlyRNAi.org-the database of the Drosophila RNAi screening center and transgenic RNAi project: 2017 update. Nucleic acids research. 45:D672–D678.

Johansson, M.J., A. Esberg, B. Huang, G.R. Bjork, and A.S. Bystrom. 2008. Eukaryotic wobble uridine modifications promote a functionally redundant decoding system. Molecular and cellular biology. 28:3301–3312.

Jumper, J., R. Evans, A. Pritzel, T. Green, M. Figurnov, O. Ronneberger, K. Tunyasuvunakool, R. Bates, A. Žídek, A. Potapenko, A. Bridgland, C. Meyer, S.A.A. Kohl, A.J. Ballard, A. Cowie, B. Romera-Paredes, S. Nikolov, R. Jain, J. Adler, T. Back, S. Petersen, D. Reiman, E. Clancy, M. Zielinski, M. Steinegger, M. Pacholska, T. Berghammer, D. Silver, O. Vinyals, A.W. Senior, K. Kavukcuoglu, P. Kohli, and D. Hassabis. 2021. Applying and improving AlphaFold at CASP14. Proteins. 89:1711–1721.

Jungfleisch, J., R. Bottcher, M. Tallo-Parra, G. Perez-Vilaro, A. Merits, E.M. Novoa, and J. Diez. 2022. CHIKV infection reprograms codon optimality to favor viral RNA translation by altering the tRNA epitranscriptome. Nature communications. 13:4725.

Kalhor, H.R., and S. Clarke. 2003. Novel methyltransferase for modified uridine residues at the wobble position of tRNA. Molecular and cellular biology. 23:9283–9292.

Kozbial, P.Z., and A.R. Mushegian. 2005. Natural history of S-adenosylmethionine-binding proteins. BMC Struct Biol. 5:19.

Lemoine, F., D. Correia, V. Lefort, O. Doppelt-Azeroual, F. Mareuil, S. Cohen-Boulakia, and O. Gascuel. 2019. NGPhylogeny.fr: new generation phylogenetic services for non-specialists. Nucleic acids research. 47:W260–w265.

Létoquart, J., N. van Tran, V. Caroline, A. Aleksandrov, N. Lazar, H. van Tilbeurgh, D. Liger, and M. Graille. 2015. Insights into molecular plasticity in protein complexes from Trm9-Trm112 tRNA modifying enzyme crystal structure. Nucleic acids research. 43:10989–11002.

Ma, N.X., B. Puls, and G. Chen. 2022. Transcriptomic analyses of NeuroD1-mediated astrocyte-to-neuron conversion. Dev Neurobiol. 82:375–391.

Maddirevula, S., S. Alameer, N. Ewida, M.M.L. de Sousa, M. Bjoras, C.B. Vagbo, and F.S. Alkuraya. 2022. Insight into ALKBH8-related intellectual developmental disability based on the first pathogenic missense variant. Hum Genet. 141:209–215.

Martin, J.L., and F.M. McMillan. 2002. SAM (dependent) I AM: the S-adenosylmethionine-dependent methyltransferase fold. Curr Opin Struct Biol. 12:783–793.

Mistry, J., S. Chuguransky, L. Williams, M. Qureshi, G.A. Salazar, E.L.L. Sonnhammer, S.C.E. Tosatto, L. Paladin, S. Raj, L.J. Richardson, R.D. Finn, and A. Bateman. 2021. Pfam: The protein families database in 2021. Nucleic acids research. 49:D412–d419.

Monies, D., C.B. Vagbo, M. Al-Owain, S. Alhomaidi, and F.S. Alkuraya. 2019. Recessive Truncating Mutations in ALKBH8 Cause Intellectual Disability and Severe Impairment of Wobble Uridine Modification. Am J Hum Genet. 104:1202–1209.

Murn, J., and Y. Shi. 2017. The winding path of protein methylation research: milestones and new frontiers. Nat Rev Mol Cell Biol. 18:517–527.

Nagarkar-Jaiswal, S., S.Z. DeLuca, P.T. Lee, W.W. Lin, H. Pan, Z. Zuo, J. Lv, A.C. Spradling, and H.J. Bellen. 2015. A genetic toolkit for tagging intronic MiMIC containing genes. eLife. 4.

Ni, J.Q., R. Zhou, B. Czech, L.P. Liu, L. Holderbaum, D. Yang-Zhou, H.S. Shim, R. Tao, D. Handler, P. Karpowicz, R. Binari, M. Booker, J. Brennecke, L.A. Perkins, G.J. Hannon, and N. Perrimon. 2011. A genome-scale shRNA resource for transgenic RNAi in Drosophila. Nature methods. 8:405–407.

Notredame, C., D.G. Higgins, and J. Heringa. 2000. T-Coffee: A novel method for fast and accurate multiple sequence alignment. J Mol Biol. 302:205–217.

Pettersen, E.F., T.D. Goddard, C.C. Huang, E.C. Meng, G.S. Couch, T.I. Croll, J.H. Morris, and T.E. Ferrin. 2021. UCSF ChimeraX: Structure visualization for researchers, educators, and developers. Protein Sci. 30:70–82.

Pieper, U., B.M. Webb, G.Q. Dong, D. Schneidman-Duhovny, H. Fan, S.J. Kim, N. Khuri, Y.G. Spill, P. Weinkam, M. Hammel, J.A. Tainer, M. Nilges, and A. Sali. 2014. ModBase, a database of annotated comparative protein structure models and associated resources. Nucleic acids research. 42:D336–346.

Saad, A.K., D. Marafi, T. Mitani, H. Du, K. Rafat, J.M. Fatih, S.N. Jhangiani, Z. Coban-Akdemir G. Baylor-Hopkins Center for Mendelian, R.A. Gibbs, D. Pehlivan, J.V. Hunter, J.E. Posey, M.S. Zaki, and J.R. Lupski. 2021. Neurodevelopmental disorder in an Egyptian family with a biallelic ALKBH8 variant. Am J Med Genet A. 185:1288–1293.

Schaffrath, R., and S.A. Leidel. 2017. Wobble uridine modifications-a reason to live, a reason to die?! RNA biology. 14:1209–1222.

Songe-Moller, L., E. van den Born, V. Leihne, C.B. Vagbo, T. Kristoffersen, H.E. Krokan, F. Kirpekar, P.O. Falnes, and A. Klungland. 2010. Mammalian ALKBH8 possesses tRNA methyltransferase activity required for the biogenesis of multiple wobble uridine modifications implicated in translational decoding. Molecular and cellular biology. 30:1814–1827.

Stoeger, T., M. Gerlach, R.I. Morimoto, and L.A. Nunes Amaral. 2018. Large-scale investigation of the reasons why potentially important genes are ignored. PLoS Biol. 16:e2006643.

Su, D., C.T. Chan, C. Gu, K.S. Lim, Y.H. Chionh, M.E. McBee, B.S. Russell, I.R. Babu, T.J. Begley, and P.C. Dedon. 2014. Quantitative analysis of ribonucleoside modifications in tRNA by HPLC-coupled mass spectrometry. Nat Protoc. 9:828–841.

Towns, W.L., and T.J. Begley. 2012. Transfer RNA methytransferases and their corresponding modifications in budding yeast and humans: activities, predications, and potential roles in human health. DNA Cell Biol. 31:434–454.

Tweedie, S., B. Braschi, K. Gray, T.E.M. Jones, R.L. Seal, B. Yates, and E.A. Bruford. 2021. Genenames.org: the HGNC and VGNC resources in 2021. Nucleic acids research. 49:D939–d946.

Uhlén, M., L. Fagerberg, B.M. Hallström, C. Lindskog, P. Oksvold, A. Mardinoglu, Å. Sivertsson, C. Kampf, E. Sjöstedt, A. Asplund, I. Olsson, K. Edlund, E. Lundberg, S. Navani, C.A. Szigyarto, J. Odeberg, D. Djureinovic, J.O. Takanen, S. Hober, T. Alm, P.H. Edqvist, H. Berling, H. Tegel, J. Mulder, J. Rockberg, P. Nilsson, J.M. Schwenk, M. Hamsten, K. von Feilitzen, M. Forsberg, L. Persson, F. Johansson, M. Zwahlen, G. von Heijne, J. Nielsen, and F. Pontén. 2015. Proteomics. Tissue-based map of the human proteome. Science. 347:1260419.

van den Born, E., C.B. Vagbo, L. Songe-Moller, V. Leihne, G.F. Lien, G. Leszczynska, A. Malkiewicz, H.E. Krokan, F. Kirpekar, A. Klungland, and P.O. Falnes. 2011. ALKBH8-mediated formation of a novel diastereomeric pair of wobble nucleosides in mammalian tRNA. Nature communications. 2:172.

van Tran, N., F.G.M. Ernst, B.R. Hawley, C. Zorbas, N. Ulryck, P. Hackert, K.E. Bohnsack, M.T. Bohnsack, S.R. Jaffrey, M. Graille, and D.L.J. Lafontaine. 2019. The human 18S rRNA m6A methyltransferase METTL5 is stabilized by TRMT112. Nucleic acids research. 47:7719–7733.

van Tran, N., L. Muller, R.L. Ross, R. Lestini, J. Letoquart, N. Ulryck, P.A. Limbach, V. de Crecy-Lagard, S. Cianferani, and M. Graille. 2018. Evolutionary insights into Trm112-methyltransferase holoenzymes involved in translation between archaea and eukaryotes. Nucleic acids research. 46:8483–8499.

Varadi, M., S. Anyango, M. Deshpande, S. Nair, C. Natassia, G. Yordanova, D. Yuan, O. Stroe, G. Wood, A. Laydon, A. Žídek, T. Green, K. Tunyasuvunakool, S. Petersen, J. Jumper, E. Clancy, R. Green, A. Vora, M. Lutfi, M. Figurnov, A. Cowie, N. Hobbs, P. Kohli, G. Kleywegt, E. Birney, D. Hassabis, and S. Velankar. 2022. AlphaFold Protein Structure Database: massively expanding the structural coverage of protein-sequence space with high-accuracy models. Nucleic acids research. 50:D439–d444.

Venken, K.J., K.L. Schulze, N.A. Haelterman, H. Pan, Y. He, M. Evans-Holm, J.W. Carlson, R.W. Levis, A.C. Spradling, R.A. Hoskins, and H.J. Bellen. 2011. MiMIC: a highly versatile transposon insertion resource for engineering Drosophila melanogaster genes. Nature methods. 8:737–743.

Wang, S., X. Liu, J. Huang, Y. Zhang, C. Sang, T. Li, and J. Yuan. 2018. Expression of KIAA1456 in lung cancer tissue and its effects on proliferation, migration and invasion of lung cancer cells. Oncology letters. 16:3791–3795.

Xu, L., X. Liu, N. Sheng, K.S. Oo, J. Liang, Y.H. Chionh, J. Xu, F. Ye, Y.G. Gao, P.C. Dedon, and X.Y. Fu. 2017. Three distinct 3-methylcytidine (m(3)C) methyltransferases modify tRNA and mRNA in mice and humans. The Journal of biological chemistry. 292:14695–14703.

Yang, W.Q., Q.P. Xiong, J.Y. Ge, H. Li, W.Y. Zhu, Y. Nie, X. Lin, D. Lv, J. Li, H. Lin, and R.J. Liu. 2021. THUMPD3-TRMT112 is a m2G methyltransferase working on a broad range of tRNA substrates. Nucleic acids research. 49:11900–11919.

Zhang, K., J.M. Lentini, C.T. Prevost, M.O. Hashem, F.S. Alkuraya, and D. Fu. 2020. An intellectual disability-associated missense variant in TRMT1 impairs tRNA modification and reconstitution of enzymatic activity. Hum Mutat. 41:600–607.

Zhou, Y., Y. Kong, W. Fan, T. Tao, Q. Xiao, N. Li, and X. Zhu. 2020. Principles of RNA methylation and their implications for biology and medicine. Biomed Pharmacother. 131:110731.

Zoghbi, H.Y., and M.F. Bear. 2012. Synaptic dysfunction in neurodevelopmental disorders associated with autism and intellectual disabilities. Cold Spring Harb Perspect Biol. 4.

