## Supplemental Figures for "Expanded tRNA methyltransferase family member TRMT9B regulates synaptic growth and function"

Figure S1

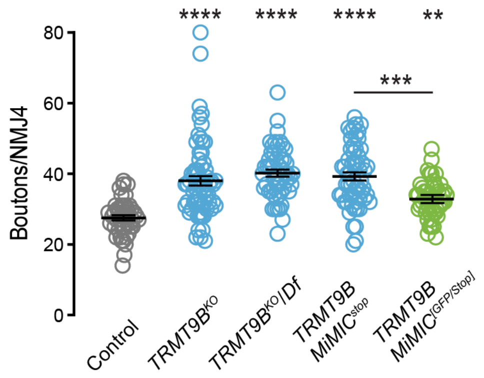


**Figure S1. TRMT9B attenuates synaptic growth.** Bouton number per NMJ in the indicated genotypes. Homozygous *TRMT9B*$^{HA+IC}$ and *TRMT9B*$^{HA+IC}$ over a deficiency exhibited similar synaptic overgrown relative to control. An independent TRMT9B MiMIC allele exhibits synaptic overgrowth that is partially rescued by conversion of one stop cassette to GFP. Kruskal-Wallis test followed by Dunn's multiple comparisons test.

Figure S2

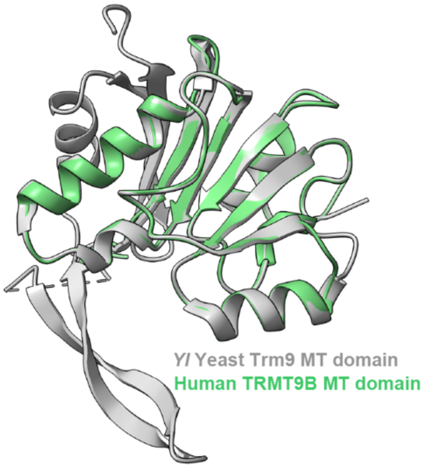


**Figure S2. Homology model of human TRMT9B and yeast Trm9**. Structural homology model of the human TRMT9B methyltransferase domain (green) with yeast Trm9 structure (gray, PBD ID: 5CM2 chain Z).

Figure S3

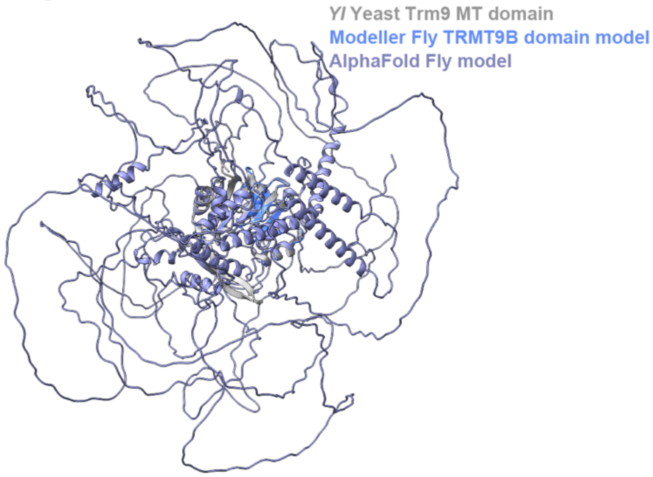


**Figure S3. Comparison of AlphaFold- and Modeller-predicted structures.** Superimposition of the predicted AlphaFold *Drosophila* TRMT9B model (purple) with the Modeller-predicted *Drosophila* TRMT9B model (blue) and yeast Trm9 structure (gray, PBD ID: 5CM2 chain Z).

Figure S4

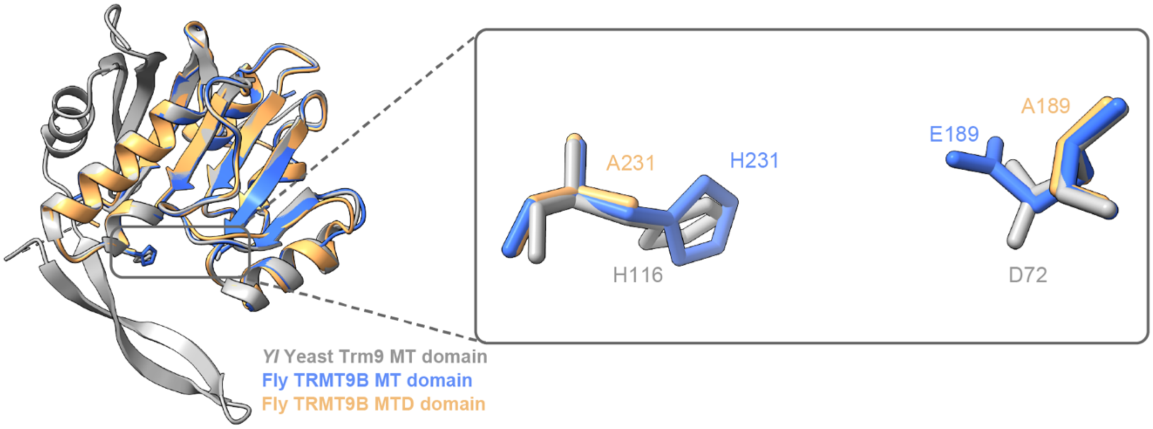


**Figure S4. Homology model of TRMT9B with methyltransferase-disrupting mutations.** Superimposition of *Drosophila* TRMT9B methyltransferase domain (blue) and methyltransferase-dead domain (MTD) models (light orange) with yeast Trm9 structure (gray, PBD ID: 5CM2 chain Z). Insert shows critical residues that were disrupted to abolish methyltransferase activity. The predicted model supports the preservation of secondary structure in TRMT9B$^{MTD}$.

Figure S5

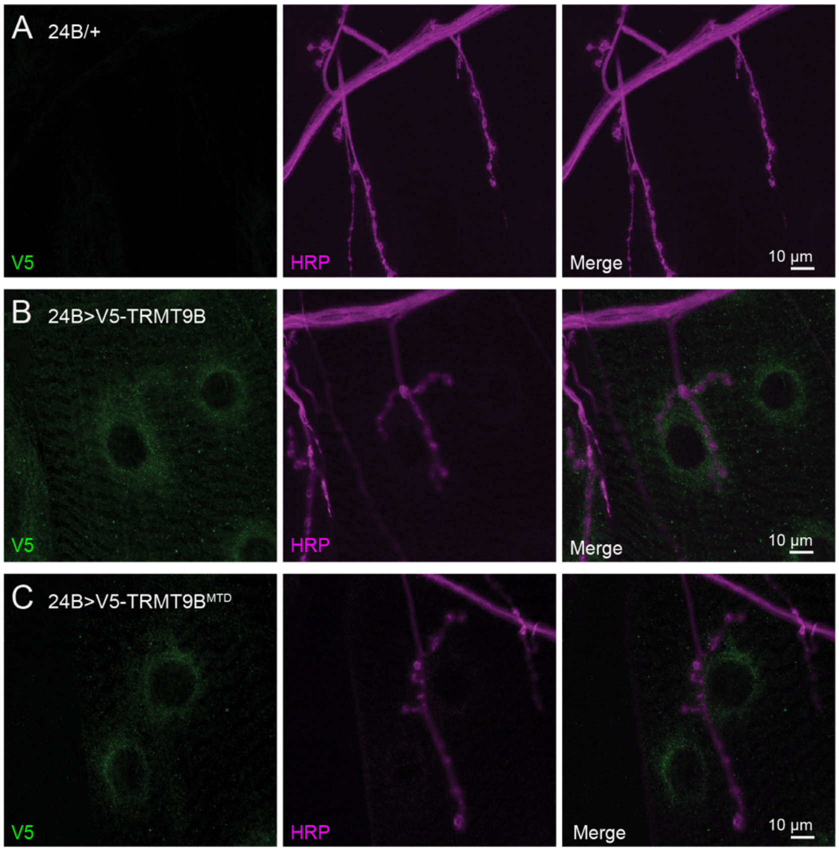


**Figure S5. wild-type and methyltransferase-dead TRMT9B transgene expression.** Confocal Z-projections of *24B-Gal4/+* control (**A**), 24B-Gal4>UAS-V5::TRMT9B (**B**) and 24B-Gal4>UAS-V5::TRMT9B$^{MTD}$ (**C**) co-labeled with antibodies against V5 (green) and the neuronal membrane marker Hrp (purple).

Figure S6

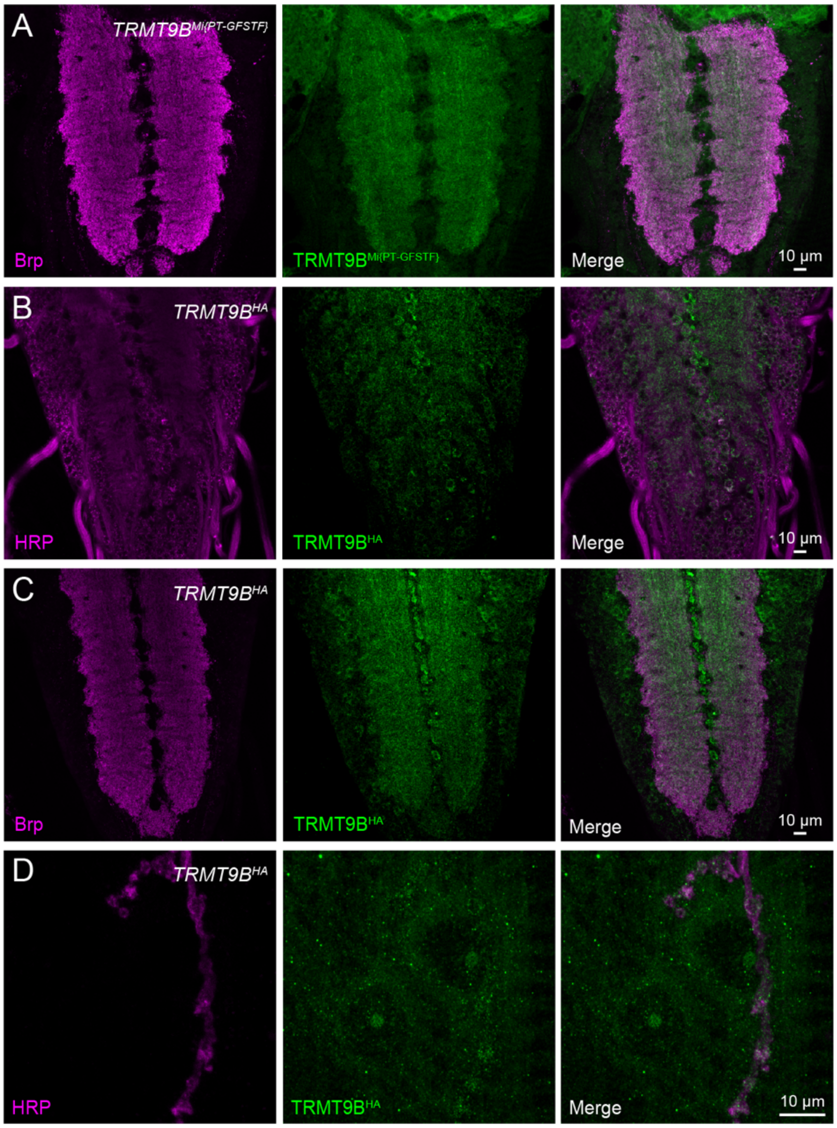


**Figure S6. TRMT9B expression in *TRMT9B*$^{MIMIC\text{-}GFP}$ and *TRMT9B*$^{HA}$.** (**A**) Confocal Z-projections of *TRMT9B*$^{MIMIC\text{-}GFP}$ (**A**) and *TRMT9B*$^{HA}$ (**B**) larval ventral ganglia co-labeled with antibodies against GFP or HA (green) and the synaptic marker Brp (purple). Confocal Z-projection of *TRMT9B*$^{HA}$ co-labeled with antibodies against HA (green) and the neuronal membrane marker HRP (purple).

Figure S7

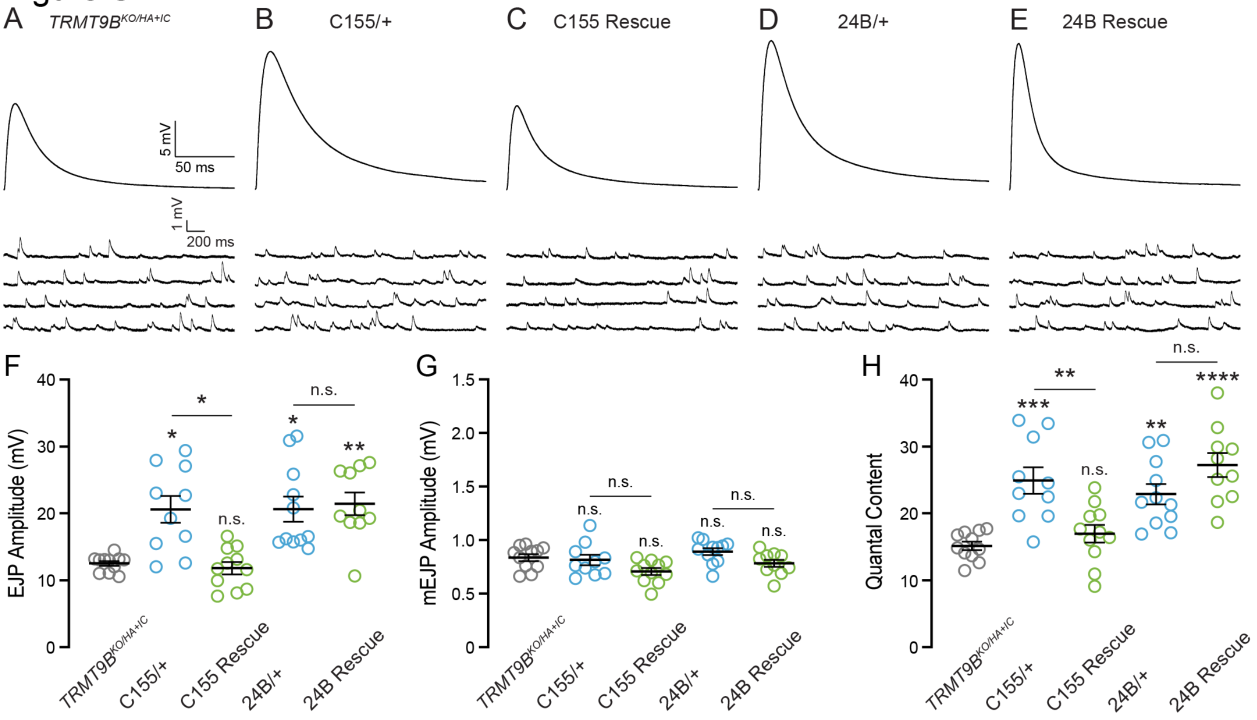


**Figure S7. Postsynaptic TRMT9B promotes neurotransmitter release.** (**A–E**) Representative traces of EJPs and mEJPs in indicated genotypes. Stimulus artifacts have been removed from representative EJPs for clarity. (**F-H**) Average EJP amplitude (**F**), mEJP amplitude (**G**) and quantal content (**H**) for the indicated genotypes. mEJP amplitude is not significantly different across larva. EJP amplitude and quantal content are significantly diminished in *TRMT9B*$^{KO/HA+IC}$ and rescued by 24B-driven, but not C155-driven, restoration of TRMT9B expression (Kruskal-Wallis non-parametric test followed by Dunn's multiple comparisons test for EJPs and ANOVA followed by Tukey's test for normally distributed mEJPs and QC). Error bars represent s.e.m.

**Supplemental files**

File S1_TRMT9B_Drosophila MT_Modeller-model_76c988b9ca53554c2f020177adf43243.pdb
File S2_TRMT9B human MT_Modeller-model_3f245a4039096aabe8e0e4bb2401c7eb.pdb
File S3_ALKBH8 Drosophila MT_Modeller-model_
3_8eaa0710e767199e6527411277bdb6cb.pdb
File S4_ALKBH8 human MT_Modeller-model_7a4c2dbde0fca4de5714657f1edb076a.pdb
File S5_TRMT9B Drosophila_MTD_Modeller-model_2b632395a5fcf3d2771385285f960342.pdb
File S6_Alphafold_TRMT9B_Drosophila_AF-Q9VBJ3-F1-model_v1.pdb
